# Apoptosis-resistance of senescent cells is an intrinsic barrier for senolysis induced by cardiac glycosides

**DOI:** 10.1101/2020.12.18.423449

**Authors:** Pavel I. Deryabin, Alla N. Shatrova, Irina I. Marakhova, Nikolay N. Nikolsky, Aleksandra V. Borodkina

**Author notes:** Correspondence to Aleksandra Borodkina, Tikhoretsky ave. 4, 194064, St-Petersburg, Russia.

## Abstract

Targeted elimination of senescent cells – senolysis – is one of the core trends in the anti-aging therapy. Cardiac glycosides were recently proved to be a broad-spectrum senolytics. Here we tested senolytic properties of cardiac glycosides towards human mesenchymal stem cells (hMSCs). Cardiac glycosides had no senolytic ability towards senescent hMSCs of various origins. Using biological and bioinformatic approaches we compared senescence development in ‘cardiac glycosides–sensitive’ A549 and ‘–insensitive’ hMSCs. The absence of senolysis was found to be mediated by the effective potassium import and increased apoptosis-resistance in senescent hMSCs. We revealed that apoptosis-resistance, previously recognized as a common characteristic of senescence, in fact, is not a general feature of senescent cells. Moreover, only apoptosis-prone senescent cells are sensitive to cardiac glycosides-induced senolysis. Thus, we can speculate that the effectiveness of senolysis might depend on whether senescent cells indeed become apoptosis-resistant compared to their proliferating counterparts.

## INTRODUCTION

Today cellular senescence is considered as a common reaction of almost all types of proliferating cells, including cancer and stem ones, as well as some post-mitotic cells to a variety of stressful stimuli (Sapieha and Mallette, 2018; Lee and Schmitt, 2019; von Zglinicki et al., 2020). The following types of cellular senescence can be distinguished accordingly to the origin of the inducing factor: replicative (due to the DNA damage at the shortened telomere ends), oncogene-induced (in response to aberrant activation of oncogenic signaling), stress-induced (mediated by the DNA damage caused by the oxidative stress, heat shock, UV and γ radiation, etc.), and chemotherapy-induced (activated in cancer cells in response to chemotherapeutic drugs) (Campisi, 1996; Serrano et al., 1997; Toussaint, et al., 2000; Roninson et al., 2001). There is a heterogeneity in markers expressed by senescent cells depending on both cell type and an insult used to induce senescence. However, there are several common features typical for the most types of senescent cells. The essential characteristic of senescence for any kind of dividing cells is the irreversible proliferation loss (Hernandez-Segura et al., 2018). The irreversibility of the cell cycle arrest is controlled by the cyclin-dependent kinase (CDK) inhibitors p16 and p21 and is often regulated by the tumor suppressor protein p53 (Hernandez-Segura et al., 2018; Wang et al., 2020). The other important features of senescent cells are the activation of a persistent DNA damage response; cell hypertrophy, which often arises as a result of impaired ribosomal biogenesis and protein synthesis; disturbance of lysosomal degradation and dysfunction of the rest degradation systems; increased activity of the specific lysosomal enzyme senescence-associated-β-galactosidase; various mitochondrial alterations; acquisition of the senescence-associated secretory phenotype, composed of pro-inflammatory factors, matrix degrading enzymes, reactive oxygen species, etc; epigenetic and chromatin landscape alterations, including formation of senescence-associated heterochromatic foci and senescence-associated distention of satellites (Nacarelli et al., 2017; Hernandez-Segura et al., 2018; Wang et al., 2020). In other words, senescent cells preserve metabolic activity and vitality, but their functioning is significantly altered comparing to the origin cells.

It is now clear that senescent cells can modify surrounding microenvironment affecting both neighboring cells and cellular niches, what may lead to tissue malfunctioning and therefore may be related to the progression of aging and age-related diseases (Munoz-Espin and Serrano, 2014; Ovadya et al., 2018). Keeping that in mind, today more attention is focused on the strategies for targeted “killing” of senescent cells (Kirkland and Tchkonia, 2020; Pignolo et al., 2020; Robbins et al., 2020). To this end, a novel class of drugs termed senolytics is actively developing. Senolytics target signaling pathways that contribute to the resistance of senescent cells towards apoptosis, thus inducing apoptosis preferentially in senescent cells (Zhu et al., 2015). The list of senolytics is constantly replenishing with the new agents. The most known compounds with the stated senolytic activity are navitoclax (Bcl-2 family inhibitor), combination of dasatinib (an inhibitor of multiple tyrosine kinases) and quercetin (a natural flavonol), Hsp90 inhibitors, MDM2 inhibitors, FOXO4-p53 interfering peptide, a BET family protein degrader, uPAR-specific CAR-T, galacto-conjugated navitoclax and various senolytic natural compounds (Zhu et al., 2015; Zhu et al., 2016; Baar et al., 2017; Fuhrmann-Strossnigg et al., 2017; Jeon et al., 2017; Zhu et al., 2017; Amor et al., 2020; Gonzalez-Gualda et al., 2020; Wakita et al., 2020). Recently, using high-throughput drug screening two independent research groups identified cardiac glycosides, particularly ouabain, digoxin and bufalin, as a broad-spectrum senolytics (Guerrero et al., 2019; Triana-Martinez et al., 2019). It is worth mentioning that almost all of the declared senolytics have limitations, such as undesirable side-effects or ineffectiveness towards some cell types.

Mesenchymal stem cells (MSCs), found virtually in all post-natal organs/tissues, are characterized by the capacity to self-renew (symmetric divisions) and to differentiate, thus contributing to maintenance and regeneration of the residing tissue (da Silva Meirelles et al., 2006). Due to these unique properties, along with the potent anti-inflammatory and immunosuppressive functions, MSCs are broadly applied in the cell replacement therapy for treatment of various diseases, including diabetes mellitus, multiple sclerosis, myocardial infarction, and so on (Zhou and Xu, 2020). However, similar to their differentiated progenies and other unipotent cells, MSCs are able to senesce either replicatively or prematurely in response to oncogenes’ activation and stressful stimuli (Burova et al., 2013; Zhou et al., 2020). MSCs’ senescence has a plenty of undesirable aftermaths, among which are exhaustion of the pool of stem cells, reduced tissue maintenance and regeneration, impaired differentiation capacity, SASP-mediated senescence spreading, reduced angiogenic properties, modification of stem cell niche (Griukova et al., 2019; Liu et al., 2020; Turinetto et al., 2016; Zhou et al., 2020). Taking into account biological significance of the proper MSCs’ functioning, senescent MSCs might be considered as the crucial target for senolysis. In line with this suggestion, a number of reviews highlighting possible advantageous outcomes of senescent MSCs removal has been published over the past year (Lee and Yu, 2020; Liu et al., 2020; Spehar et al., 2020; Zhou et al., 2020). Nevertheless, there are only few experimental data regarding this issue (Grezella et al., 2018; Peng et al., 2020; Sharma et al., 2020; Zhang et al., 2020).

Within the present study we demonstrate for the first time that cardiac glycosides, namely ouabain and bufalin, fail to display senolytic activity towards human MSCs (hMSCs) derived from endometrium (END-MSCs), adipose tissue (AD-MSCs), dental pulp (DP-MSCs) and Warton jelly (WJ-MSCs). Additionally, we confirm that both cardiac glycosides are able to induce apoptosis preferentially in senescent A549, as it was previously described in the pilot studies (Guerrero et al., 2019; Triana-Martinez et al., 2019). By assessing alterations in ionic homeostasis caused by the Na^+^/K^+^-ATPase blocking and expression levels of the related genes we reveal that the absence of ouabain-induced senolysis might be mediated by the enhanced effectiveness of the compensatory K^+^ import in senescent END-MSCs compared to senescent A549. Furthermore, using advanced bioinformatics we demonstrate that senescence of END-MSCs, resistant to ouabain-induced senolysis, is accompanied by the acquisition of apoptosis-resistant phenotype, while senescence of ouabain-sensitive A549 is not. Therefore, we provide clear evidence that apoptosis-resistance is not a general feature of senescent cells. Based on the data obtained we conclude that the effectiveness of senolytic approaches might depend on whether cells indeed became apoptosis-resistant during senescence development.

## RESULTS

### 1. H_2_O_2_ –treated END-MSCs enter the premature senescence

Within the present study we used human mesenchymal stem cells isolated from desquamated endometrium (END-MSCs), which satisfy the minimum criteria suggested by the ISCT for defining hMSCs (Deryabin et al., 2019). Previously, we have developed the reliable experimental model to study various aspects of the premature senescence of END-MSCs (Burova et al., 2013). Namely, we have shown that END-MSCs subjected to sublethal oxidative stress gradually acquired all the typical features of senescent cells, including persistent DNA damage foci and active DNA damage response, irreversible cell cycle arrest mediated by the classical p53/p21/Rb pathway, proliferation loss, cell hypertrophy, appearance of SA-β-Gal staining and development of senescence-associated secretory phenotype (Burova et al., 2013; Borodkina et al., 2014; Griukova et al., 2018). Here we applied the designed model to study senolytic effect of ouabain as well as to reveal the underlying intracellular alterations in ion homeostasis. Initially, by estimating the most common parameters we confirmed that single-dose (1 h, 200 μM) H_2_O_2_ treatment was sufficient to induce senescence in END-MSCs. Importantly, stressed END-MSCs were considered senescent in two weeks after the oxidative stress; therefore, all the senescence markers were assessed not earlier than 14 days after H_2_O_2_ treatment. Indeed, H_2_O_2_-treatment of END-MSCs led to proliferation block, cell hypertrophy as indicated by the increased cell size, accumulation of the lipofuscine detected by the elevation of autofluorescence, appearance of the senescence-associated β-galactosidase staining (SA-β-Gal), activation of the p21/Rb pathway, and loss of HMGB1 together supporting senescence establishment under the chosen experimental conditions (Figure 1A–C,E,F). The most harmful effects of senescent cells at tissue and organismal levels are believed to be the consequence of their prolonged vitality. In line with this point, H_2_O_2_-treated END-MSCs retained high viability even at the late stages of senescence development (viability of senescent END-MSCs was assessed in 17 days (14 days + 3 days) after the initial oxidative stress) (Figure 1D). To sum up, sublethal H_2_O_2_-treatment of END-MSCs is the appropriate model of the premature senescence and is relevant to investigate the effects of senolytic compounds on END-MSCs.

**Figure 1.**
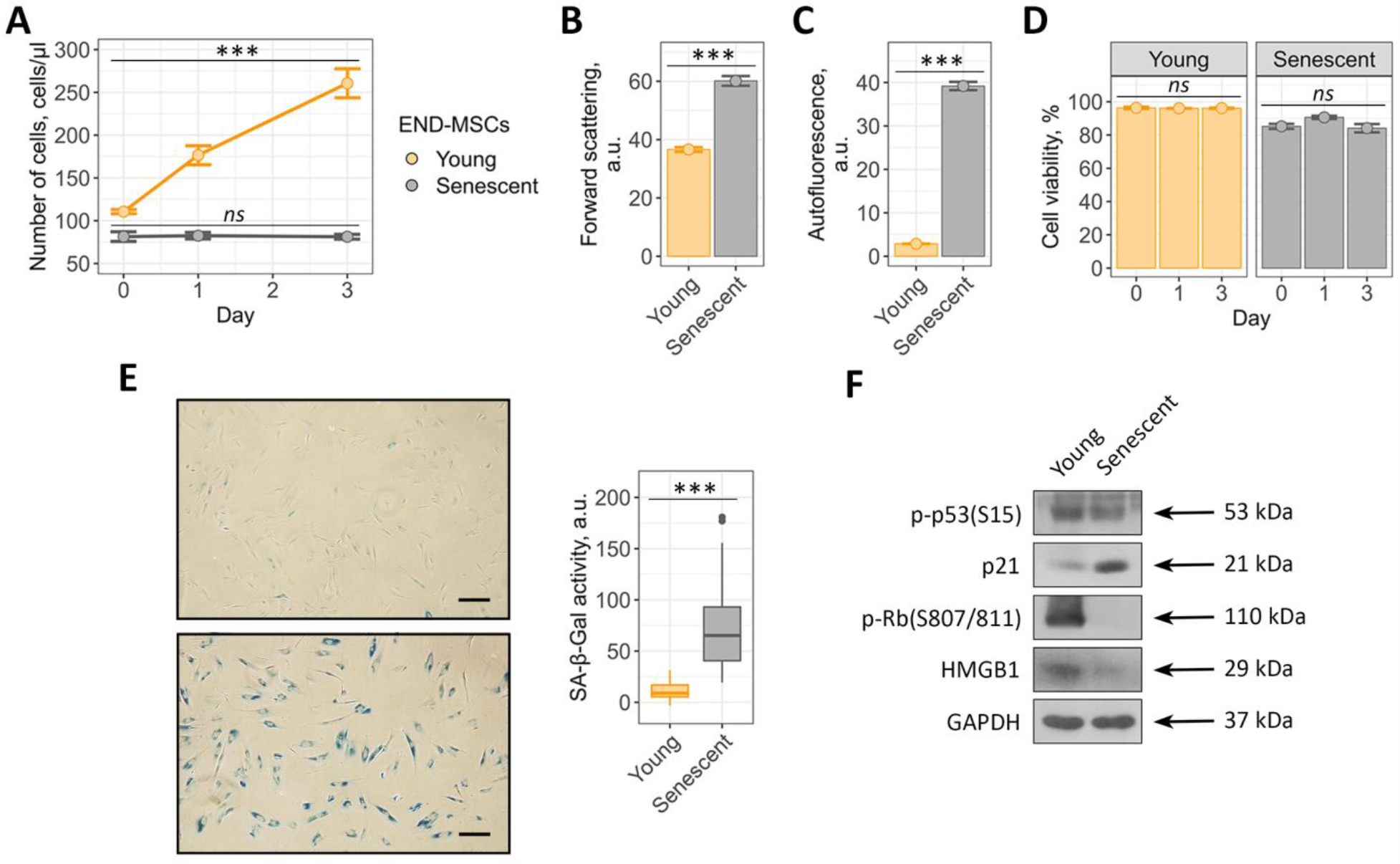
Validation of oxidative stress induced END-MSCs premature senescence model. Senescent END-MSCs (**A**) lose proliferation, (**B**) undergo hypertrophy, (**C**) acquire elevated autofluorescence, retain high cell viability (**D**) and (**E**) display SA-β-Gal activity compared to the young ones. (**F**) Phosphorylation levels of p53 and Rb and expression levels of p21 and HMGB1 proteins in young and senescent END-MSCs. Values presented are mean ± s.d. For multiple groups comparisons at (**A**) and (**D**) one-way ANOVA was applied, n = 3, ns – not significant, *** p < 0.001. For pair comparisons at (**B**), (**C**), and (**E**) Welch’s t-test was used, n = 3 for **b** and **c**, n = 50 for **e**, *** p < 0.001. Scale bars for images are 500 μm. GAPDH was used as loading control.

### 2. Cardiac glycoside ouabain has no senolytic activity towards senescent END-MSCs in a wide concentration range

Recent evidence suggest that cardiac glycosides, including ouabain, digoxin, bufalin, represent a family of compounds with senolytic activity (Guerrero et al., 2019; Triana-Martinez et al., 2019). Even though today cardiac glycosides are considered as the broad-spectrum senolytics, the data regarding their effects towards senescent hMSCs are lacking. Therefore, here we tested whether ouabain has the potential to induce death selectively in senescent END-MSCs. To do so, we assessed viability of young and senescent END-MSCs after treatment with ouabain at the wide concentration range (from 10^−7^ to 10^−5^ M). Interestingly, neither concentration applied led to the noticeable decrease in the viability of both young and senescent cells on the first day after ouabain application (Figure2–figure supplement 1). However, on the third day after ouabain treatment we revealed significant dose-dependent decline in the viability of young END-MSCs (Figure 2A, Figure2–figure supplement 1). Unexpectedly, senescent cells turned out to be more resistant towards ouabain at each concentration tested (Figure 2A, Figure2–figure supplement 1). In line with this result, 10^−6^ M ouabain led to more prone apoptotic death in young cells (20.36 % An+/PI- and 34.99 % An+/PI+) compared to senescent ones (15.69 % An+/PI- and 12.42 % An+/PI+) (Figure 2B). Obtained results clearly demonstrate that in context of senescent END-MSCs ouabain has no senolytic activity.

**Figure 2.**
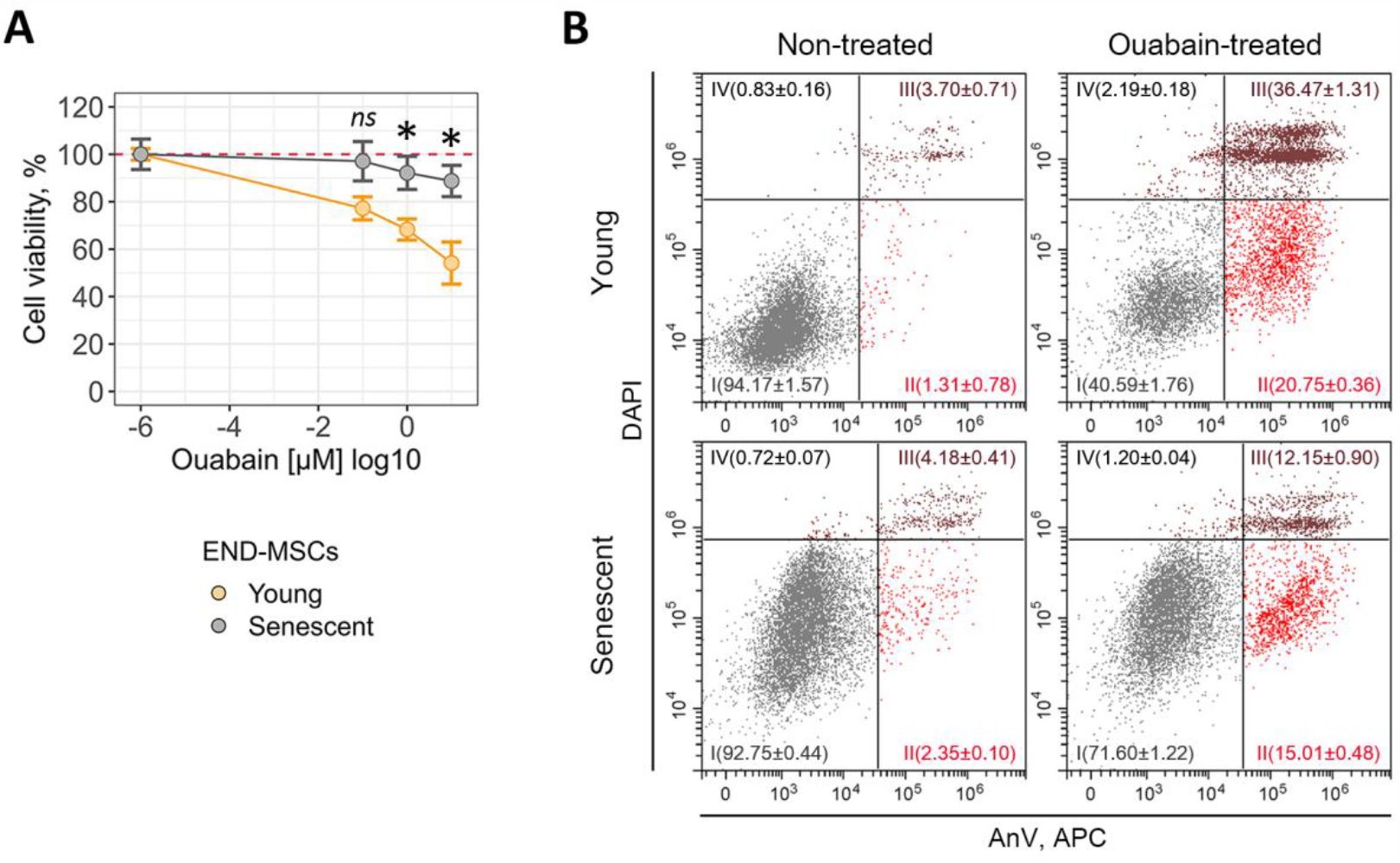
Ouabain has no senolytic activity towards senescent END-MSCs in a wide concentration range. Relative cell viability (%) of young and senescent END-MSCs in 3 days after treatment with 10^−7^, 10^−6^, 10^−5^ M ouabain. (**B**) Apoptosis induction in END-MSCs upon 10^−6^ M ouabain assessed by Annexin V/DAPI double staining. n = 3 independent experiments. All data correspond to the mean ± s.d. Statistical significance was assessed by the Welch’s t-test: ns – not significant, ***p < 0.05.

### 3. Ouabain is able to induce apoptosis selectively in senescent A549 cells

Taking into account the fact that our results do not correspond with the recently published evidences regarding broad-spectrum senolytic action of cardiac glycosides, we decided to reproduce this effect using cellular model described in the relevant studies (Guerrero et al., 2019; Triana-Martinez et al., 2019). Thus, we performed series of experiments using young and senescent A549 lung carcinoma cells. Senescence in A549 cells was induced by etoposide treatment. Etoposide-induced senescence of A549 cells is a frequently used and thus well characterized model of therapy-induced senescence (Wang et al., 2017). Also, ouabain was shown to selectively kill etoposide-treated senescent A549 cells (Guerrero et al., 2019). To prove senescence in A549, we assessed proliferation rate, cell size, accumulation of lipofucine, SA-β-Gal staining, activation status of the p53/p21/Rb pathway and expression level of HMGB1 (Figure 3A–E).

**Figure 3.**
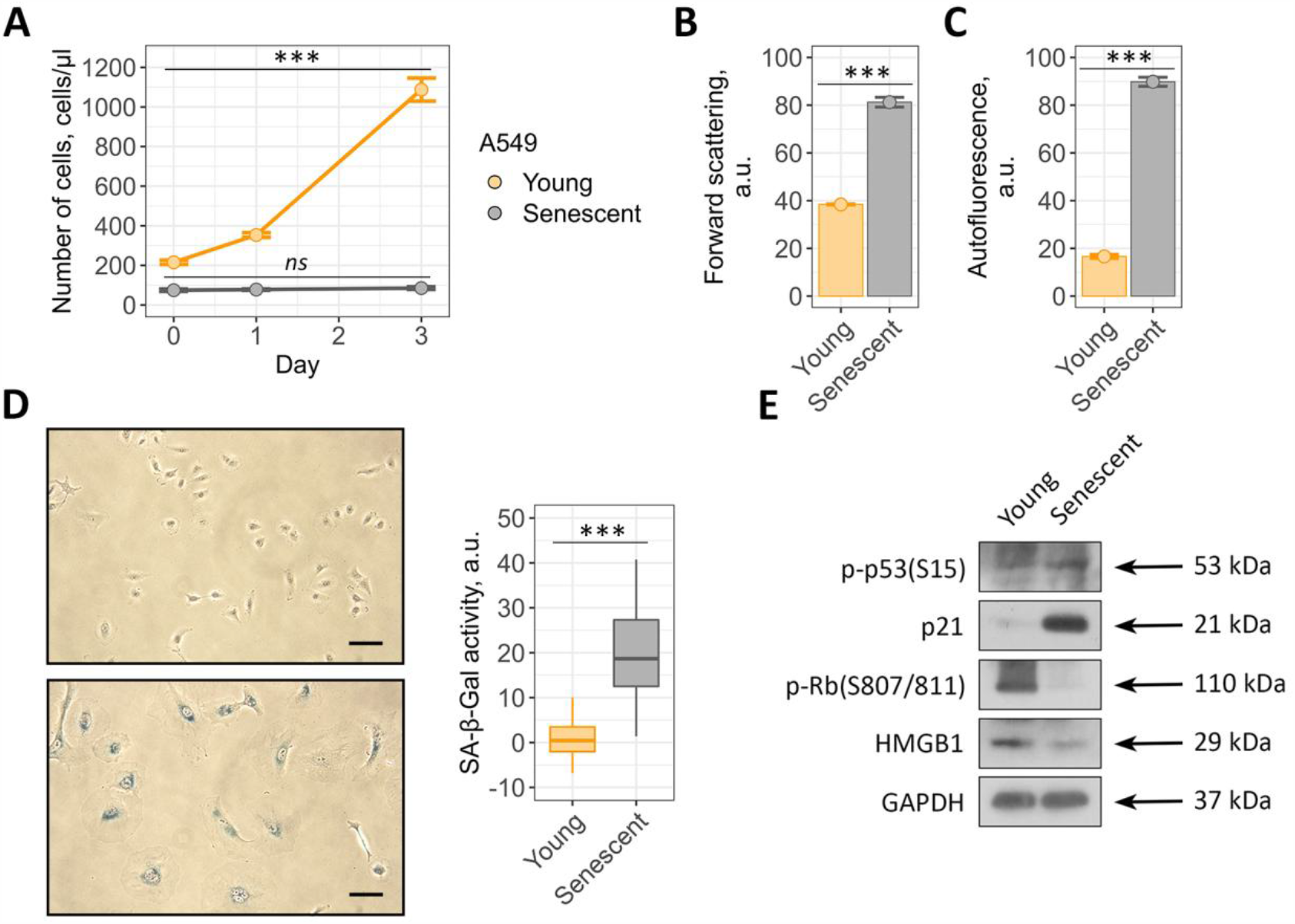
Validation of etoposide induced A549 senescence model. Senescent A549 cells display (**A**) loss of proliferation, (**B**) acquire elevated cell size, (**C**) autofluorescence level and (**D**) SA-b-Gal activity compared to the young ones. (**E**) Phosphorylation levels of p53 and Rb and expression levels of p21 and HMGB1 proteins in young and senescent A549. Values presented are mean ± s.d. For multiple groups comparisons at (**A**) one-way ANOVA was applied, n = 3, ns – not significant, *** p < 0.001. For pair comparisons at (**B**), (**C**), and (**D**) Welch’s t-test was used, n = 3 for (**B**) and (**C**), n = 50 for (**D**), *** p < 0.001. Scale bars for images are 500 μm. GAPDH was used as loading control.

To verify senolytic activity of ouabain towards senescent cancer cells, we first estimated dose-dependent cell viability. In line with our results described above, we were not able to detect any significant decline in the number of viable young or senescent A549 cells within 24 h after ouabain application (Figure 4––figure supplement 1). However, in 3 days after treatment ouabain significantly reduced viability of senescent A549 cells in a dose-dependent manner, while the number of young A549 cells decreased to a much lesser extent (Figure 4A, Figure 4––figure supplement 1). Namely, approximately 90 % of young cells preserved viability at 10^−6^ M ouabain compared to 50 % of senescent A549 cells treated with the same dose (Figure 4A).

**Figure 4.**
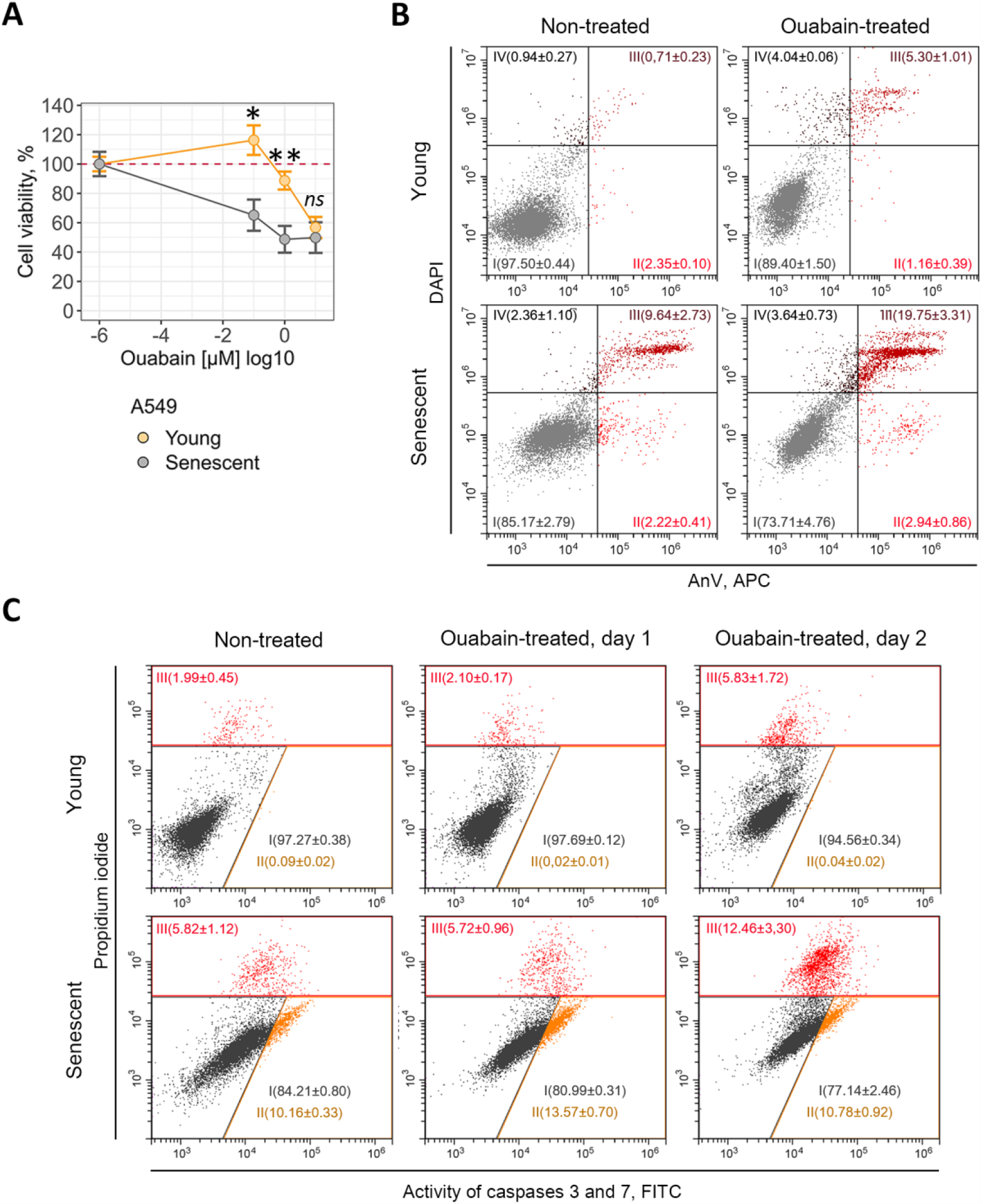
Ouabain induces caspase-dependent apoptosis selectively in senescent A549 cells. (**A**) Relative cell viability (%) of young and senescent A549 in 3 days after treatment with 10^−7^, 10^−6^, 10^−5^ M ouabain. (**B**) Apoptosis induction in A549 upon 10^−6^ M ouabain assessed by Annexin V/DAPI double staining. (**C**) Activity of Caspase-3/7 estimated in young and senescent A549 treated with 10^−6^ M ouabain. n = 3 independent experiments. All data correspond to the mean ± s.d. Statistical significance was assessed by the Welch’s t-test: * ns – not significant, p < 0.05, **p < 0.01.

According to the published data, ouabain triggered caspase-3-dependent apoptosis in senescent A549 cells (Triana-Martinez et al., 2019). Indeed, we revealed significant increase in the double positive AnV/PI-fraction in senescent cancer cells treated with 10^−6^ M ouabain (Figure 4B). Furthermore, using fluorescent assay we observed activation of caspase-3 in ouabain-treated senescent A549 cells (Figure 4C). Taken together, these results are completely consistent with the data described by other authors, and confirm senolytic activity of ouabain towards A549.

### 4. Cardiac glycoside bufalin is able to induce senolysis in A549 cells, while fails to kill senescent END-MSCs

To verify the opposite senolytic action of cardiac glycosides towards A549 and END-MSCs, we applied bufalin, another compound with the stated senolytic activity belonging to the cardiac glycosides family (Guerrero et al., 2019). Bufalin had almost no effect on the viability of young and senescent END-MSCs in a wide concentration range (from 10^−7^ to 10^−5^ M) (Figure 5A, Figure 5–figure supplement 1A). Contrarily, both types of A549 cells were sensitive to bufalin (Figure 5B, Figure 5–figure supplement 1B). Moreover, senescent A549 cells were more prone towards bufalin-induced cell death than their young counterparts (Figure 5B). The results described above confirm that cardiac glycosides indeed have senolytic activity towards A549, but these compounds turned to be ineffective for targeted death induction in senescent END-MSCs.

**Figure 5.**
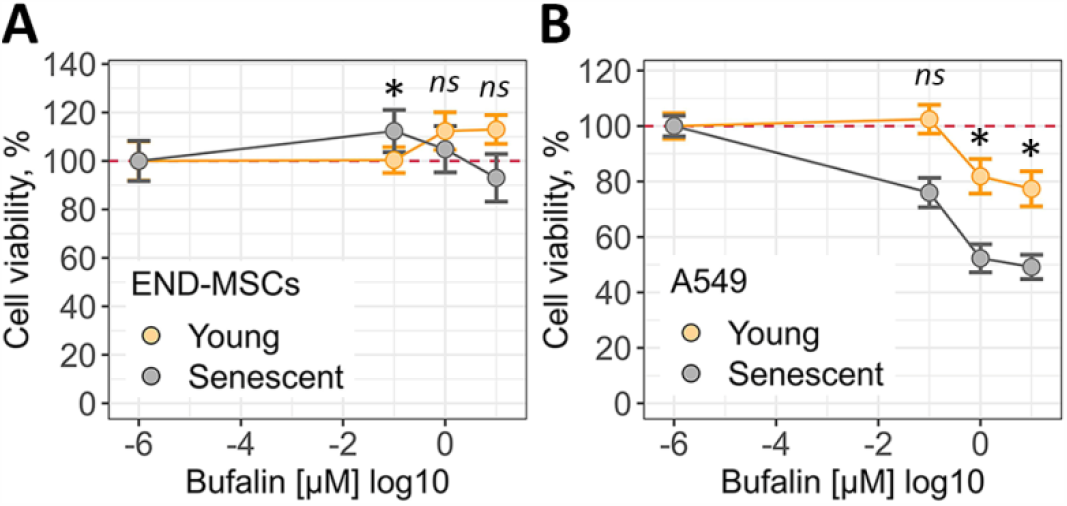
Cardiac glycoside bufalin is able to induce senolysis in A549 cells and fails to kill senescent END-MSCs. (**A**) Relative cell viability (%) of young and senescent END-MSCs in 3 days after treatment with 10^−7^, 10^−6^, 10^−5^ M bufalin. (**B**) Relative cell viability (%) of young and senescent A549 in 3 days after treatment with 10^−7^, 10^−6^, 10^−5^ M bufalin. All data correspond to the mean ± s.d. Statistical significance was assessed by the Welch’s t-test: ns – not significant, *p < 0.05.

### 5. Ouabain has no senolytic action towards human mesenchymal stem cells of various origins

To broaden our observations regarding the absence of ouabain-induced senolysis in END-MSCs, we analyzed ouabain effects on hMSCs isolated from other sources including adipose tissue, dental pulp and Wharton’s jelly. To additionally strengthen our data, we applied different models of senescence – replicative senescence for AD-MSCs, doxorubicine-induced senescence for DP-MSCs and oxidative stress-induced senescence for WJ-MSCs (Figure 6A–F).

**Figure 6.**
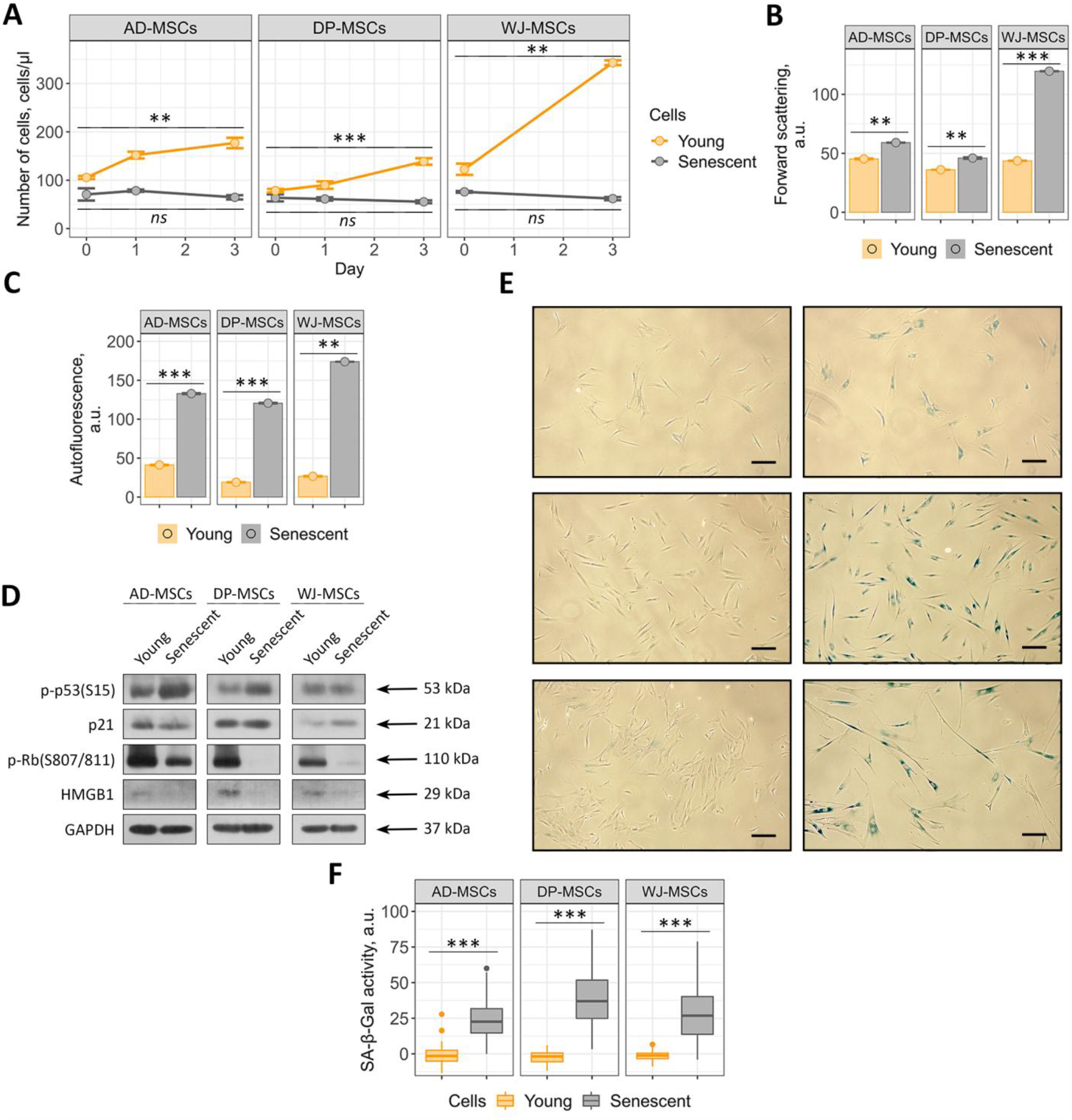
Validation of the three additional hMSCs senescence models, replicative for AD-MSCs and premature for DP-MSCs (doxorubicin induced) and WJ-MSCs (oxidative stress induced). (**A**) Loss of proliferation ability and acquisition of elevated (**B**) cell size, (**C**) autofluorescence levels and (**E**) and (**F**) SA-β-Gal activity by senescent hMSCs compared to the young cells. (**D**) Phosphorylation levels of p53 and Rb and expression levels of p21 and HMGB1 proteins in young and senescent hMSCs. Values are mean ± s.d. For multiple groups comparisons at a one-way ANOVA was applied, n = 3, ns – not significant, ** p < 0.01, *** p < 0.001. For pair comparisons at (**A**), (**B**), (**C**), and (**E**) Welch’s t-test was used, n = 3 for (**A**), (**B**), (**C**) n = 50 for (**F**), ** p < 0.01, *** p < 0.001. Scale bars for images are 500 μm. GAPDH was used as loading control.

As shown in Figure 7A–C, various types of hMSCs differed in the viability upon ouabain treatment, for example both young and senescent DP-MSCs were much more resistant to ouabain action than WJ-MSCs (Figure 7B,C, Figure 7––figure supplement 1B,C). Nevertheless, ouabain was not able to induce senolysis in either type of hMSCs (Figure 7A–C, Figure 7–– figure supplement 1A–C). Together, the obtained data demonstrate that the absence of cardiac glycoside-induced senolysis is a common feature for various types of hMSCs.

**Figure 7.**
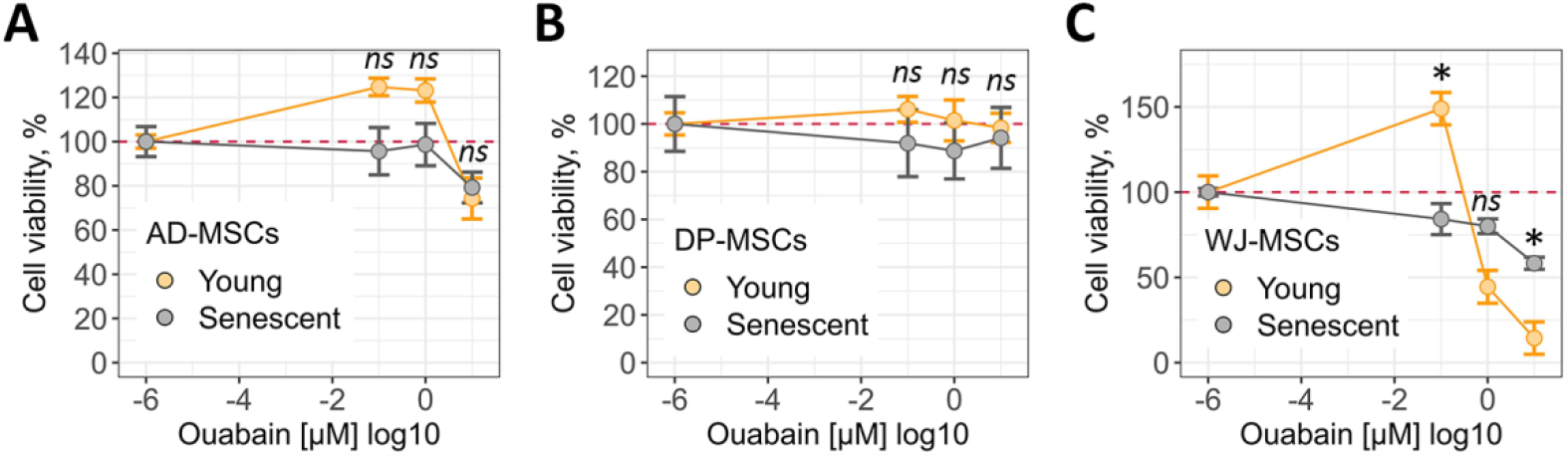
Ouabain has no senolytic action towards AD-MSCs, DP-MSCs and WJ-MSCs. (**A**), (**B**), (**C**) Relative cell viability (%) of young and senescent AD-MSCs, DP-MSCs and WJ-MSCs after treatment with 10^−7^, 10^−6^, 10^−5^ M ouabain, respectively. All data correspond to the mean ± s.d. Statistical significance was assessed by the Welch’s t-test: ns – not significant, *p < 0.05, **p < 0.01.

### 6. Disturbance of K^+^/Na^+^ homeostasis caused by ouabain leads to different physiological reactions in young and senescent A549 and END-MSCs

Having established the fact that senolytic effect of ouabain is cell type-dependent, further we tried to elucidate what may underlie the revealed differences in responses of hMSCs and A549. The main molecular mechanism of action of various cardiac glycosides is inhibition of Na^+^/K^+^-ATPase (Skou, 1988). Na^+^/K^+^-ATPase is a plasma membrane enzyme that pumps sodium out of the cell while pumping potassium into the cell against concentration gradients, and thus participating in maintenance K^+^ and Na^+^ homeostasis. Therefore, we asked whether there might be any differences in basal and ouabain-induced ion homeostasis between young and senescent cells of two types. We first estimated intracellular contents of K^+^ and Na^+^ in young and senescent END-MSCs and A549 cells in normal culture conditions and upon ouabain application. Of note, within the present study the measured ions were normalized per total protein in the same cells to quantify intracellular ion content. Young and senescent cells of both types displayed typical ion distribution, namely high intracellular K^+^ and low intracellular Na^+^ (Figure 8A,B). As expected, ouabain shifted intracellular ion contents, leading to the vice versa intracellular ionic ratios in both cell lines (Figure 8A,B). Importantly, the overall tendency of ionic changes induced by ouabain was similar for END-MSCs and A549, indicating the absence of any significant variations in the primary response of both cell types to ouabain.

**Figure 8.**
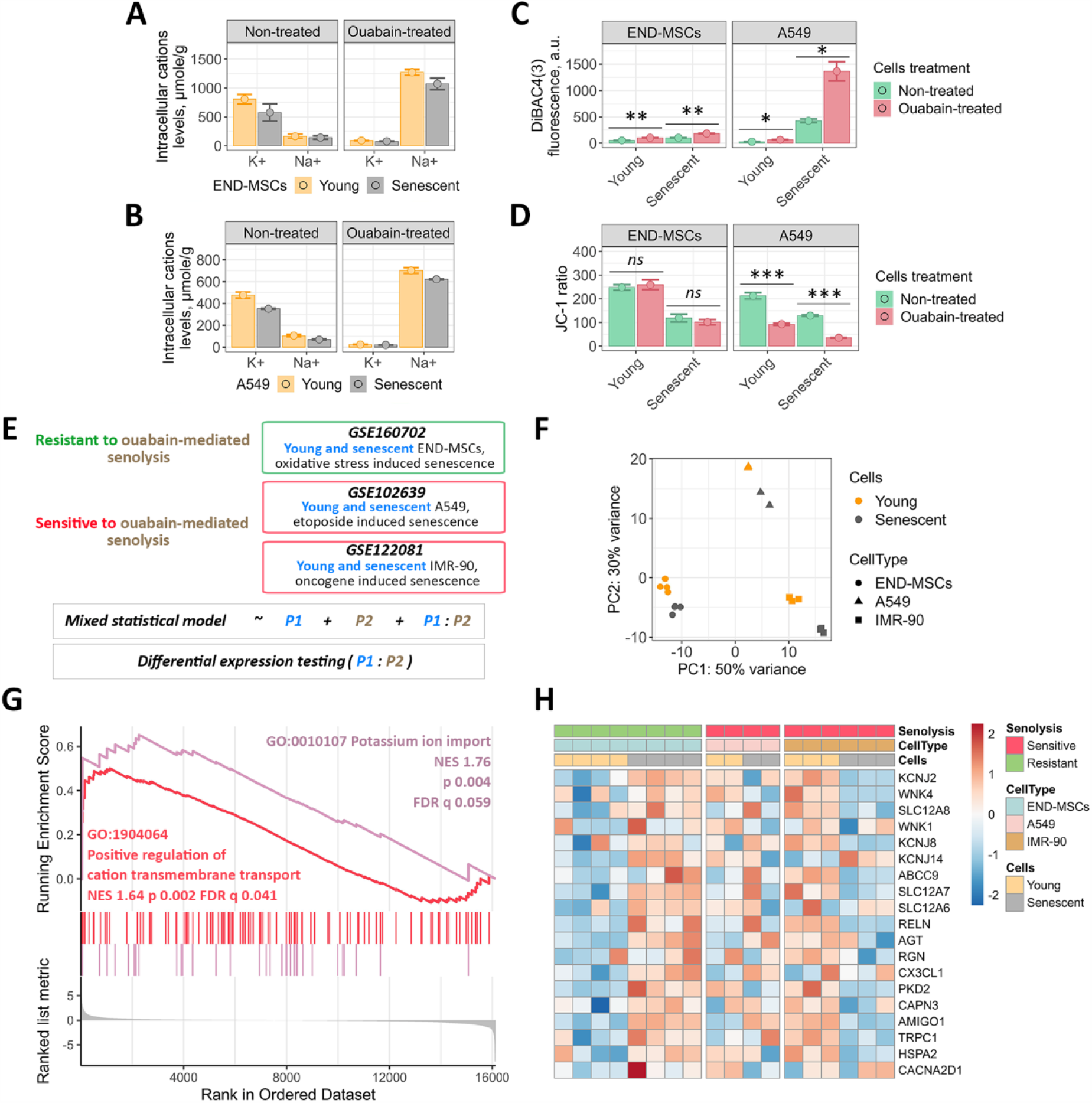
Ouabain-resistant senescent END-MSCs are able to restore disturbance of K^+^/Na^+^ homeostasis caused by ouabain via effective K^+^ import, while ouabain-sensitive senescent cells lack this ability. (**A**), (**B**) Basal and ouabain-modified intracellular levels of K^+^ and Na^+^ in young and senescent END-MSCs and A549, respectively. (**C**) Membrane potential determination of young and senescent END-MSCs and A549 cells before and 24 h after ouabain treatment, using fluorescent probe DiBAC4(3). (**D**) Mitochondrial membrane potential evaluation of young and senescent END-MSCs and A549 cells before and 24 h after ouabain treatment, using fluorescent probe JC-1. (**E**) Design of the comparative bioinformatic analysis of three independent RNA-Seq datasets. (**F**) Principal Component Analysis for the subset of genes related to the GO term “cellular senescence” (GO 0090398) based on the combined data from three independent RNA-Seq datasets. (**G**) GSEA results for the differentially expressed genes between the cells resistant to ouabain-mediated senolysis (END-MSCs) and ones sensitive to ouabain-mediated senolysis (A549 and IMR-90) for ‘Positive regulation of cation transmembrane transport’ and ‘Potassium ion import’ biological processes. (**H**) Heatmap reflecting the core enrichment genes for ‘Positive regulation of cation transmembrane transport’ and ‘Potassium ion import’ biological processes. Values are mean ± s.d. Statistical significance was assessed by the two-tailed Student’s t-test: ns – not significant, *p < 0.05, **p < 0.01, ***p < 0.001.

The disturbance in K^+^ and Na^+^ homeostasis caused by ouabain should ultimately lead to the dissipation of the plasma membrane potential. Thus, further we used specific fluorescent probe DiBAC4(3) to measure transmembrane potential of non-excitable cells. Increased fluorescence of this dye reflects membrane depolarization. In each cell type tested ouabain induced predictable membrane depolarization, confirming ionic disbalance (Figure 8C). Notably, we observed comparable levels of membrane depolarization in young and senescent END-MSCs, while senescent A549 characterized by significantly more depolarized membrane compared to young A549 cells (Figure 8C).

Other important consequences of ionic deregulation are changes in mitochondrial membrane potential (MMP). Thus, using fluorescent probe JC-1 we assessed MMP in young and senescent cells upon ouabain treatment. It should be specifically highlighted that senescent cells are commonly characterized by decreased MMP reflecting malfunctioning of mitochondria, alterations in energy metabolism and increased intracellular reactive oxygen species levels (Passos et al., 2010). As expected, MMP in senescent END-MSCs and A549 cells was lower than in the young ones, though this decrease did not affect viability of senescent cells (Figure 1D, Figure 3A). Similar to the above results, ouabain induced pronounced MMP depolarization in senescent A549 compared to their proliferating counterparts, while effect on MMP of young and senescent END-MSCs was minimal (Figure 8D).

Summarizing the results described within this part, we can conclude that blocking Na^+^/K^+^-ATPase by ouabain causes comparable alterations in intracellular ions in young and senescent cells of both types, however, in case of senescent A549, contrarily to senescent END-MSCs, these alterations lead to the dramatic depolarization of plasma membrane and drop of MMP.

### 7. Alterations in the mechanisms of K^+^ import mediate ouabain-resistance of senescent END-MSCs

To clarify mechanisms mediating ouabain-induced changes in membrane polarization and MMP between senescent A549 and END-MSCs, further we employed advanced bioinformatic approach. The aim of the analysis was to reveal possible transcriptomic features mediating cell type dependent ouabain-resistance or -sensitivity of senescent cells. In other words, we asked the question: how senescence of END-MSCs (ouabain-resistant cells) differs from senescence of A549 (ouabaine-sensitive cells)? To answer this question, we utilized three independent RNA sequencing (RNA-Seq) datasets. The first one was RNA-Seq analysis of young and senescent END-MSCs performed by us. The second one was the dataset for young and senescent A549 downloaded from GEO (Wang et al., 2017). Of note, senescence inducing conditions coincided with those applied in the present study and in the pilot studies of cardiac glycosides-induced senolysis. And the third dataset was for young and senescent IMR-90 obtained directly from the study on cardiac glycosides as senolytic compounds (Guerrero et al., 2019). Importantly, IMR-90 were proved to be prone for ouabaine-induced senolysis. It should be noted that comparing only two datasets relevant for END-MSCs and A549 would ultimately lead to improper results, as in such analysis the sought-for difference between senescence process in END-MSCs and A549 would be indistinguishable from the variations mediated by diverse technical batch effects (i.e. type of sequencing machine, the conditions of the sample ran). To minimize any batch effects, we compared one dataset for ouabain-resistant cells (END-MSCs) and two for ouabain-sensitive cells (A549, IMR-90). To validate the relevance of the further comparative analysis, we performed a Principal Component Analysis for the subset of genes related to the GO term “cellular senescence” (GO 0090398) based on the summary dataset containing samples from all three described experiments. Indeed, for each cell type samples clustered into two separate groups – the young and the senescent ones (Figure 8F).

To perform analysis of differentially expressed genes (DEGs) during senescence of ouabain-resistant and -sensitive cells, we generated the statistical model that summed up two predictors and their interaction. The schematic presentation of the analysis is displayed in Fig. 8e. In brief, the first predictor in the model divided all samples by young and senescence subgroups, the second – by ouabain-resistantce or ouabain-sensitivity, and the last component in the model reflected the interaction of both predictors. The result of differential expression testing for the last variable is presented in Supplementary table 1. Importantly, there was no significant difference in the expression levels of Na^+^/K^+^-ATPase subunits during senescence of ouabain-resistant or -sensitive cells (Supplementary Table 2). This result correlated well with the similarity in intracellular K^+^ and Na^+^ shifts upon Na^+^/K^+^-ATPase inhibition described in the previous part (Figure 8A,B).

We then conducted gene set enrichment analysis (GSEA) in Gene Onthology (GO) terms for Biological Processes (BP) (Supplementary Table 3). Notably, among the significantly enriched processes we found ‘Potassium ion import’ and ‘Positive regulation of cation transmembrane transport’ to be up-regulated during senescence of ouabain-resistant END-MSCs compared to senescence of ouabain-sensitive cells (Figure 8G). The core enrichment gene lists related to these processes included *KCNJ2, KCNJ14, KCNJ8, SLC12A7, SLC12A8, WNK4* which were significantly upregulated in senescent END-MSCs (Figure 8H). The proteins encoded by *KCNJ* genes are inward-rectifier type potassium channels, which have a greater tendency to allow potassium to flow into a cell rather than out of a cell. SLC12 is a family of cation-coupled chloride transporters. *WNK4* gene encodes for serine/threonine kinase that acts as an activator of sodium-coupled chloride cotransporters and inhibitor of potassium-coupled chloride cotransporters.

Together, these findings allowed suggesting that senescent END-MSCs can restore depleted intracellular K^+^ levels more effectively than senescent A549 cells and thus can manage with ouabain-induced ionic disbalance.

### 8. Senescent END-MSCs display elevated apoptosis-resistance compared to the young ones, while senescent A549 do not

Our data demonstrating the difference in the maintenance of K^+^ homeostasis in senescent END-MSCs and A549 cells together with the established role of intracellular K^+^ in the regulation of cell death prompted us to compare the overall stress resistance acquired during senescence of both cell types. It is well known that exogenous stresses are accompanied by the disturbance in ionic homeostasis, leading to a rapid exchange of various ions, including K^+^ between the cell and its environment (Kondratskyi et al., 2015). The outcomes of stressful influences are largely dependent on the ability of cells to restore the appropriate intracellular ionic balance. As shown in Figure 8G,H, senescence of END-MSCs was accompanied by the upregulation of K^+^ import suggesting that during senescence development END-MSCs enhance their ability to restore K^+^ levels. Interestingly, we were not able to reveal similar tendency for A549 senescence, thus, A549 seemed not to acquire any specific features related to the K^+^ import during senescence progression. According to the literary data, stress-induced decrease in intracellular K^+^ and inability to restore its level favor activation of caspases and nucleases and thus are proposed to be a pro-apoptotic factors (Kondratskyi et al., 2015).

In line with this suggestion, using bioinformatic analysis of the above RNA-seq datasets we revealed significant down-regulation of the processes related to apoptosis during senescence of END-MSCs compared to A549 and IMR-90 (Figure 9A). Moreover, senescence of A549 and IMR-90 characterized by significant up-regulation of pro-apoptotic genes, including *BAD, BAX, BOK, BAK-1, NOXA* (Figure 9B). These data indicate that END-MSCs acquire apoptosis-resistant phenotype during senescence, while senescent IMR-90 and A549 became apoptosis-prone (Figure 9B).

**Figure 9.**
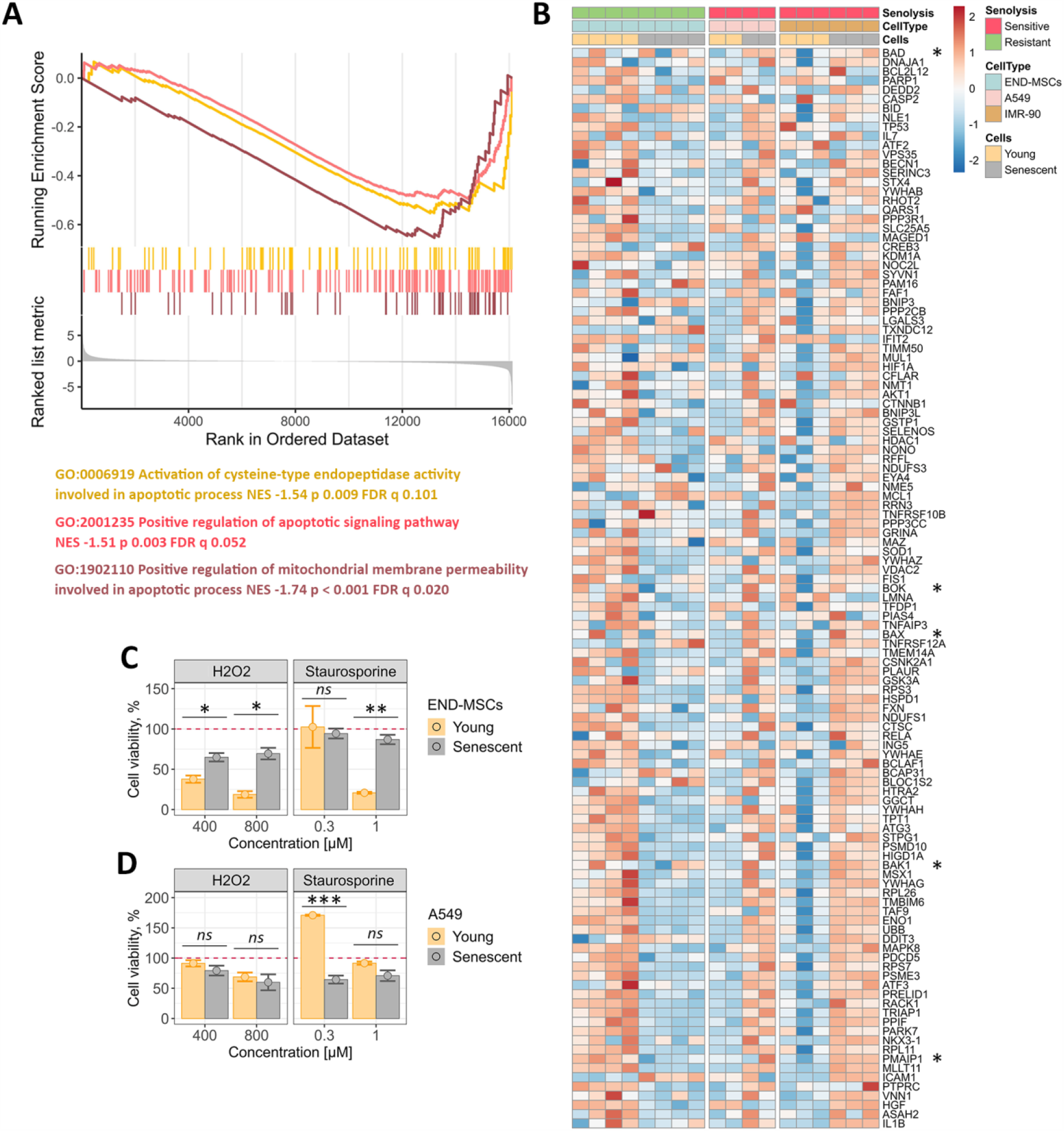
Senescence of ouabain-resistant END-MSCs is accompanied by acquisition of apoptosis-resistant phenotype, while ouabain-sensitive cells become apoptosis-prone during senescence. (**A**) GSEA results for the DEGs between the cells resistant to ouabain-mediated senolysis (END-MSCs) and the ones sensitive to ouabain-mediated senolysis (A549 and IMR-90) in the apoptosis-related terms. (**B**) Heatmap reflecting expression of the core enrichment genes for apoptosis-related terms from **a**. The common pro-apoptotic genes are marked with asterisks. (**C**), (**D**) Relative cell viability (%) of young and senescent END-MSCs or A549 treated either with H_2_O_2_ or staurosporine at indicated concentrations. Values are mean ± s.d. Statistical significance was assessed by the two-tailed Student’s t-test: *p < 0.05, **p < 0.01, ***p < 0.001, ns – not significant.

We next tested whether the observed distinctions in apoptosis background during senescence of END-MSCs and A549 may have an impact on the stress resistance of senescent cells. To do so, we assessed cell viability upon oxidative stress and staurosporine, typically applied to induce apoptosis. Indeed, senescent END-MSCs turned out to be significantly more resistant towards both stresses compared to their young counterparts (Figure 9C). Contrarily, A549 cells demonstrated vice versa situation, as senescent cells displayed tendency to be more sensitive towards stress-induced cell death (Figure 9D). This observation shifts the existing paradigm that enhanced death resistance is a common feature of any kind of senescent cells, demonstrating its obvious cell specificity.

Several crucial conclusions and the outflowing assumptions can be done based on the data presented within this study. Cardiac glycosides have no senolytic effects on hMSCs of different origin. The absence of this effect is probably mediated by the upregulation of the systems supporting ionic balance in this type of senescent cells. Other less evident reason for ouabain-resistance of hMSCs might be connected with the increase in overall resistance to stress-induced death during their senescence. Senescent A549 cells are ouabain-sensitive, however, this type of senescent cells is less resistant to cell death induced by either stress compared to the young ones. We can speculate that preferential killing of senescent cells by senolytics might work for those types of senescent cells that in fact did not become more resistant to cell death (Figure 10).

**Figure 10.**
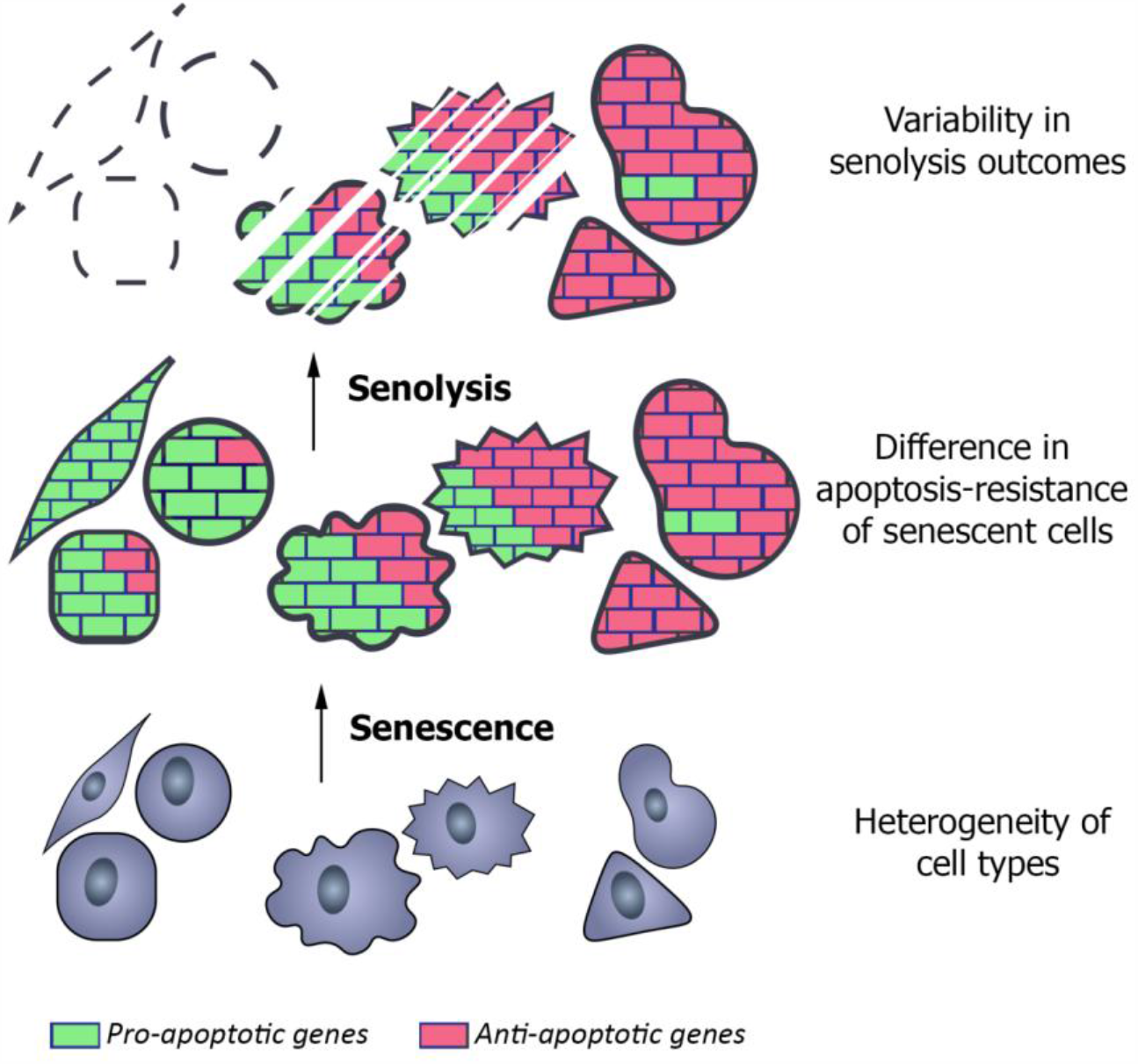
Proposed model suggesting apoptosis-resistance of senescent cells to be an intrinsic barrier for senolysis induced by cardiac glycosides. Becoming senescent heterogeneous cell types acquire apoptosis-prone, apoptosis-resistant or mixed phenotypes. Senolytics, e.g. cardiac glycosides, can effectively eliminate only apoptosis-prone senescent cells.

## DISCUSSION

Within the present study we tested senolytic effects of cardiac glycosides, namely ouabain and bufalin, towards hMSCs. Both compounds were recently identified to have selective cytotoxic effect on senescent cells (oncogene-/therapy-induced senescence models) of different origins, including primary cells IMR-90, HUVEC, ARPE-19, T/C-28, and cancer cells SKHep1, A549, SK-Mel-5, MCF7, HCT116, HaCat, H1299, U373-MG, H1755 (Guerrero et al., 2019; Triana-Martinez et al., 2019). Unexpectedly, we were not able to reveal any preferential cytotoxic effects neither of ouabain nor of bufalin on senescent END-MSCs compared to the young ones. Importantly, the absence of senolysis was verified using hMSCs obtained from adipose tissue, dental pulp and Warton jelly, and besides various senescence models. The latter led us to the suggestion that resistance to cardiac glycoside-induced senolysis might be the common feature for hMSCs. At the same time we were able to induce apoptosis preferentially in senescent A549, thus reproducing senolytic effect of cardiac glycosides described in the relevant articles (Guerrero et al., 2019; Triana-Martinez et al., 2019). With cells resistant and sensitive to ouabain-induced senolysis (ouabain-resistant/ouabain-sensitive) available, we tried to elucidate fundamental causes underlying this difference.

The common molecular mechanism of cardiac glycosides’ action is binding to the Na^+^/K^+^-ATPase and blocking its activity. By hydrolyzing ATP this enzyme ensures pumping Na^+^ out of the cell and importing K^+^ into the cells, and thus maintains physiological electrochemical gradient, ionic homeostasis, cellular pH and cell volume that are essential for cell survival and functioning (Chen et al., 2014). Therefore, we speculated that the senolytic ability of ouabain might depend on the severity of K^+^/Na^+^ disbalance in the treated cells. However, we did not reveal any significant differences in cationic shifts between ouabain-resistant and -sensitive models, as the dynamics of ouabain-induced alterations in intracellular K^+^ and Na^+^ were similar between both types of young and senescent cells. This similarity can be partially explained by the absence of any significant alterations in the expression levels of Na^+^/K^+^-ATPase subunits during senescence of ouabain-resistant or -sensitive cells as indicated by the bioinformatic analysis of the RNA-seq data.

Generally speaking, the impact of Na^+^/K^+^-ATPase inhibiting on cell survival was shown to be cell-type specific. For example, ouabain was shown to potentiate apoptosis in lymphocytes, Jurkat cells, canine epithelial cells, while it failed to induce death in epithelial cells from rat aorta, rat cerebral granule cells and porcine renal proximal tubular LLC-PK1 lymphocytes despite similar modulation of the cationic ratio (Ledbetter et al., 1986; Olej et al., 1998; Isaev et al., 2000; Bortner et al., 2001; Orlov et al., 2001; Zhou et al., 2001; Pchejetski et al., 2003). One of the proposed mechanisms of proapoptotic ouabain action is depletion of intracellular K^+^ that favors apoptotic shrinkage, activation of caspases and initiation of apoptotic programming (Kondratskyi et al., 2015; Chen et al., 2014). Considering comparable K^+^ loss but opposite effect on death induction obtained for ouabain-resistant and ouabain-sensitive models, we suggested the differences in cation’s compensatory systems. Drop of intracellular K^+^ level should essentially lead to the activation of the K^+^ import from the extracellular space to compensate the lack of this cation. Therefore, the effectiveness of K^+^ restoration might underlie apoptosis predisposition upon ouabain treatment. To test this suggestion, we applied complex bioinformatic analysis comparing alterations in transcriptomic signatures during senescence of ouabain-resistant cells (END-MSCs) and ouabain-sensitive cells (A549 and IMR-90). We observed strong upregulation of the processes related to potassium ion import during END-MSCs senescence, while during senescence of ouabain-sensitive A549 and IMR-90 these processes remained unchanged or even decreased. Based on that we can speculate that senescent END-MSCs can effectively cope with ouabain-induced K^+^ depletion via active cation importing systems, what prevents apoptosis induction. Contrarily, ouabain-sensitive senescent A549 and IMR-90 seem to be less able to overcome K^+^ loss, thus are apoptosis-prone. In favor of this assumption, ouabain induced more significant plasma and mitochondrial membranes depolarization in senescent A549 cells, demonstrating pronounced ionic disbalance typical for dying cells.

Rather than being the unique feature of ouabain-induced apoptosis, drop of intracellular K^+^ content is probably the common characteristic of apoptotic death triggered by various stresses, e.g. staurosporine treatment and oxidative stress (Kondratskyi et al., 2015). Accordingly, decreased ability of senescent A549 to restore cytoplasmic K^+^ should ultimately lead to decline in the overall stress resistance. However, this notion contradicts with the modern definition of cell senescence, which states that senescent cells are highly resistant to apoptosis (Zhu et al., 2015). It should be specifically highlighted that enhanced apoptosis resistance formed the basis for senolytics development (Zhu et al., 2015; Zhu et al., 2016).

The first mention of the increased stress resistance of senescent cells dates back to 1995, when replicatively senescent fibroblasts were found to be more resistant to serum withdrawal compared to the young ones (Wang, 1995). This initial study was followed by the limited number of investigations indicating that senescent cells are more resistant to UV-light, staurosporine, thapsigargin and other stresses than their proliferating counterparts (Marcotte et al., 2004; Ryu et al., 2007). However, we are far from the first to raise the question of the enhanced apoptosis-resistance as the general feature of senescent cells. Thus, in the review published in 2003 it was reported “… apoptosis resistance is not a general feature of senescent cells, which may also be apoptotic prone depending on cell type and apoptotic stimuli…” (Soti et al., 2003). Indeed, several data demonstrated higher sensitivity of senescent cells to stress-induced apoptosis than of young cells (Hoffmann et al., 2001; Hampel et al., 2004; Jeon and Boo, 2013). For example, senescent HUVECs were more prone to apoptosis induced by oxidized LDL or TNFα compared to young cells (Hoffmann et al., 2001). Despite of the obvious controversy, the last doubt appeared in 2015 and sounded as follows: “It seems doubtful that global apoptosis resistance is a general feature of senescence cells” (Burton and Faragher, 2015). In the same year the first study uncovering senolysis was published (Zhu et al., 2015). Within this study the authors highlighted increased expression of pro-survival gene networks in senescent cells that contributed to their enhanced resistance to apoptosis. Thus, initially senolytics represented the drugs to target proteins that protected senescent cells from apoptosis. The discovery of senolytics together with the impressive results of senescent cells’ removal using INK-ATTAC transgenic mice triggered a flurry of studies searching for the new compounds with senolytic activity (Zhu et al., 2015; Baker et al., 2016; Zhu et al., 2016; Baar et al., 2017; Fuhrmann-Strossnigg et al., 2017; Jeon et al., 2017; Zhu et al., 2017; Guerrero et al., 2019; Triana-Martinez et al., 2019; Amor et al., 2020; Gonzalez-Gualda et al., 2020; Wakita et al., 2020). Over the last two years a plenty of reviews on senolytics was published (Paez-Ribes et al., 2019; Soto-Gamez et al., 2019; Kirkland and Tchkonia, 2020; Pignolo et al., 2020; Robbins et al., 2020; Wang et al., 2020). However, since 2015 enhanced apoptosis resistance of senescent cells was not the matter of debate, though, in fact, it was not tested within these studies.

Interestingly, when comparing transcriptomic signatures of ouabain-resistant cells (END-MSCs) and ouabain-sensitive cells (A549 and IMR-90), we found that only senescence of END-MSCs was accompanied by acquisition of noticeable anti-apoptotic profile. On the contrary, senescence of A549 and IMR-90 characterized by significant up-regulation of pro-apoptotic genes, including *BAD, BAX, BOK, BAK-1, NOXA* and so on. On the one hand, these results provide additional molecular explanation for the absence of ouabain-induced senolysis in END-MSCs, and, on the other hand, clearly demonstrate that A459 and IMR-90 become predisposed to apoptosis during senescence. Of note, our data regarding transcriptomic alterations in genes related to apoptosis in senescent IMR-90 coincided with the results presented, yet not discussed, in the article published by Guerrero et al., as the datasets for young and senescent IMR-90 cells originated from this study (Guerrero et al., 2019). The precise analysis of the heatmap provided within this study revealed upregulation of *BAX, BID, BAD, BAK1* and *NOXA* in senescent IMR-90 compared to the young cells. Furthermore, Baar et al. also revealed “unexpected” upregulation of proapoptotic and downregulation of antiapoptotic genes in senescent IMR-90, mentioning that senescent IMR-90 should be primed to undergo apoptosis (Baar et al., 2017).

In order to strengthen our observation, we compared resistance of young and senescent END-MSCs and A459 to the most common apoptosis-inducing stimuli – oxidative stress and staurosporine. According to the results of our bioinformatic analysis, senescent END-MSCs turned out to be far more stress-resistant than young cells. At the same time, senescent A549 demonstrated the opposite reaction being slightly more susceptible than young cells to both types of stressful stimuli. Therefore, senolytic action of cardiac glycosides, at least for A549, might be mediated by the predisposition of senescent cells of this origin to apoptosis compared to their young counterparts, whereas the absence of ouabain-induced senolysis in hMSCs might be explained by the increase in overall apoptotic resistance during senescence of these cells. If the latter is true, then senescent hMSCs are expected to be resistant towards other senolytic compounds. Indeed, by analyzing the existing evidence we found that various compounds with the stated senolytic activity, including navitoclax, nicotinamide riboside, danazol geldanamycin, ganetespib, fisetin, BCL-XL inhibitors, quercetin turned to be ineffective in the targeted removal of senescent human preadipocytes (fibroblast like precursor cells derived from human adipose tissue) and hMSC from bone marrow (Zhu et al., 2016; Fuhrmann-Stroissnigg et al., 2017; Zhu et al., 2017; Grezella et al., 2018).

The data regarding the absence of senolysis in hMSCs somewhat contradict with the inspiring results obtained using in vivo mice models, demonstrating positive outcomes upon systemic application of senolytics (Peng et al., 2020; Zhang et al., 2020). For example, it was shown that clearance of senescent salivary gland stem cells using ABT263 may prevent radiotherapy-induced xerostomia (Peng et al., 2020). Furthermore, removal of senescent bone marrow MSCs by quercetin improved bone marrow formation (Zhang et al., 2020). Importantly, it was shown that senescent mouse and rat MSCs unlike hMSCs are responsive to senolytics. For example, quercetin, quercetin + dasatinib, ABT263 significantly removed senescent mouse MSCs (Zhu et al., 205). Also, 17-DMAG has been shown to greatly reduce senescent bmMSCs in a progeroid mouse model (Fuhrmann-Stroissnigg et al., 2017). Therefore, the improvements in various organs and systems functioning described within these studies are speculated to be the consequence of the targeted removal of senescent mouse stem cells. However, these senolytics have not yet shown any functional rejuvenation of hMSCs. Such an obvious distinction between the responsiveness of mice and human MSCs towards senolysis suggests that improvements from senolytic therapies may not be conserved across the species. The results obtained for senolytic compound UBX0101 can be considered as a good illustration confirming the last statement. This agent has been shown to clear senescent cells in a mouse model of osteoarthritis, which significantly reduced pain and promoted repair of the damaged cartilage (Jeon et al., 2017). Unfortunately, UBX0101 failed to attenuate disease progression and pain in patients with moderate-to-serve painful osteoarthritis during phase II clinical trial (NCT04129944; Dolgin, 2020; Garth, 2020). The similar concerns are raised about senolytic combination dasatinib+quercetin (Garth, 2020).

To sum up, two important conclusions flow out from the obtained data. The former demonstrated that cardiac glycosides are unable to clear senescent hMSC. The absence of senolysis might be mediated by effective K^+^ cellular import and increased apoptosis resistance in senescent hMSCs. The latter is more fundamental and reveals that apoptosis resistance is not a general feature of senescent cells, as during senescence some cells acquire ‘apoptosis-resistant’ phenotype, while others do not. Importantly, only apoptosis-prone senescent A549 cells could be effectively cleared by cardiac glycosides. Based on that we can speculate that the effectiveness of other senolytic approaches might depend on whether senescent cells are indeed apoptosis-resistant compared to their proliferating counterparts. Finally, though ‘senolytics’ is a hot topic and a lot of inspiring data are published, conclusions regarding the effectiveness of senolysis should be taken with caution as heterogeneity of cell senescence still remains a ‘puzzle’.

## METHODS

### Cells

END-MSCs were previously isolated from desquamated endometrium (Burova et al., 2013). The study was reviewed and approved by the Local Bioethics Committee of the Institute of Cytology of the Russian Academy of Sciences. Written informed consent was obtained from all patients who provided tissue. WJ-MSCs, DP-MSCs, AD-MSCs and A549 were obtained from the Russian Collection of Cell Cultures (Institute of Cytology, Saint-Petersburg, Russia). A549 line was authenticated by karyology, tumorigenicity, isoenzyme (LDH and G6PD) tests, and STR analysis. Cells were cultured in complete medium DMEM/F12 (Gibco BRL) supplemented with 10 % FBS (HyClone), 1 % penicillin-streptomycin (Gibco BRL) and 1 % glutamax (Gibco BRL). All cells were routinely tested for mycoplasma contamination.

### Senescence- and stress-inducing conditions

For oxidative stress-induced senescence END-MSCs/WJ-MSCs were treated with 200 µM/100 µM H_2_O_2_ (Sigma) for 1 h. For doxorubicine-induced senescence DP-MSCs were treated with 100 nM of doxorubicine (Veropharm) for 3 days. In each case cells were considered senescent not earlier then 14 days after treatment. For replicative senescence AD-MSCs earlier than 4^th^ passage were identified as young cells and later than 10^th^ as senescent ones. For chemotherapy-induced senescence A549 were treated with 2 µM etoposide (Veropharm) for 3 days and analyzed not earlier than 7 days after senescence induction. In this case, treatment design completely coincided with those described in the study from which RNA-seq data originated (Wang et al., 2017). Two cardiac glycosides we applied as senolytic compounds – Ouabain (Sigma), Bufalin (Calbiochem).

In order to compare stress-resistance between young and senescent END-MSCs or A549, cells were treated either with 400/800 µM H_2_O_2_ (Sigma) for 1 h or with 0.3/1 µM staurosporine (Sigma). Cell viability was analyzed in 48 h after stress induction.

### Flow cytometry analysis

Measurements of cell viability, proliferation, cell size, autofluorescence, apoptosis rates, caspase 3/7 cleavage, mitochondrial membrane potential, membrane depolarization were carried out by flow cytometry. Flow cytometry was performed using the CytoFLEX (Beckman Coulter) and the obtained data were analyzed using CytExpert software version 2.0. Adherent cells were rinsed twice with PBS and harvested by trypsinization. Detached cells were pooled and resuspended in fresh medium and then counted and analyzed for autofluorescence. In order to access cell viability, 50 μg/ml propidium iodide (Life Technologies) was added to each sample just before analysis. The cell size was evaluated by cytometric forward light scattering of PI-negative cells. Apoptosis induction was verified using Annexin-V-APC (Invitrogen) and DAPI (Sigma) co-staining following manufactures instructions. Caspase activity was assessed using CellEvent(tm) Caspase-3/7 Green Flow Cytometry Assay Kit (Invitrogen) following manufactures protocol. Loss of mitochondrial membrane potential was assessed using the ratiometric dye JC-1 (Invitrogen). The staining procedure was carried out in accordance with the manufacture’s protocol. For membrane depolarization we used DiBAC4(3) fluorescent probe (Invitrogen). Gating strategy for flow cytometry analysis is provided in Supplementary Figure 1.

### Senescence-associated β-Galactosidase staining

Senescence-associated β-Galactosidase staining was performed using senescence β-galactosidase staining kit (Cell Signaling Technology) according to manufacturer’s instructions. The kit detects β-galactosidase activity at pH 6.0 in cultured cells that is present only in senescent cells and is not found in pre-senescent, quiescent or immortal cells. Quantitative analysis of images was produced with the application of MatLab package, according to the algorithm described in the relevant paper (Shlush et al., 2011). For each experimental point not less 50 randomly selected cells were analyzed.

### Western blotting

Western blotting was performed as described previously^41^. SDS-PAGE electrophoresis, transfer to nitrocellulose membrane and immunoblotting with ECL (Thermo Scientific) detection were performed according to standard manufacturer’s protocols (Bio-Rad Laboratories). Antibodies against the following proteins were used: glyceraldehyde-3-phosphate dehydrogenase (GAPDH) (clone 14C10), phospho-p53 (Ser15), p21 (clone 12D1), phospho-Rb (Ser807/811), HMGB1 (clone D3E5), as well as horseradish peroxidase-conjugated goat anti-rabbit IgG. All antibodies were purchased from Cell Signaling.

### Determination of cell ions

Cellular K^+^ and Na^+^ contents were measured by emission photometry in an air–propane flame using a Perkin-Elmer AA 306 spectrophotometer as described previously (Marakhova et al., 1998). In summary, the cells were pelleted in RPMI medium, washed five times with MgCl2 solution (96 mM) and treated with 5% trichloroacetic acid (TCA). TCA extracts were analyzed for ion content. Intracellular K+, Na+, and Rb+ contents were determined by flame emission on a Perkin-Elmer AA 306 spectrophotometer. TCA precipitates were dissolved in 0.1 N NaOH and analyzed for protein by Lowry procedure. The cell ion content was expressed as amount of ions per amount of protein in each sample analyzed.

### RNA-Seq analysis

Samples from the following three Gene Expression Omnibus datasets were used in the analysis: GSE102639 (GSM2742113 - GSM2742114 and GSM2742121 - GSM2742122 for young and senescent A549 cells, accordingly); GSE122081 (GSM3454482 - GSM3454484 and GSM3454500 - GSM3454502 for young and senescent IMR-90 cells, accordingly); and our dataset GSE160702 (GSM4877895 - GSM4877898 and GSM4877907 - GSM4877910 for young and senescent END-MSCs, respectively). Data for all the datasets were processed in the same way.

Raw reads data underwent quality filtering and adapter trimming via FilterByTile and BBDuk scripts from the BBtools package (version 38.75) using the default options. The remaining reads were additionally filtered and trimmed with the use of trimFilter script from the FastqPuri package (version 1.0.7). Trimming operation was applied for both ends of reads if they contained N’s or their quality was below the quality threshold set to 27, all reads shorter than 25 bases were discarded. The quality control of trimming was held with the FastQC software (version 0.11.7) and FastqPuri scripts. The reads, having passed all operations, comprised no less than 90% of the initial data.

Transcript abundances were estimated using the Salmon lightweight-mapping (version 1.1.0) running in the selective alignment mode. The list of decoys was generated based on the Gencode human reference genome GRCh38.p13 (release 33) and used further for building the index on concatenated transcriptome and genome Gencode reference files (release 33) using k-mer size of 21. Mapping operations were run with additional flags --numBootstraps 30 --seqBias --gcBias --validateMappings. Resulting mapping rates were around 70%.

Further data processing was performed using R version 3.6.3 with the Tidyverse collection of packages (version 1.3.0). Estimated gene counts, metadata and transcript ranges were loaded into R and summarized to a gene level using tximeta (version 1.4.5). Resulting count matrix was filtered to contain rows having at least 5 estimated counts across all samples, the resulting matrix contained 20400 genes.

For the PCA and heatmap representation the read counts were normalized using rlog transformation from the DESeq2 package (version 1.26.0). Heatmaps were constructed with the use of genefilter (version 1.38.0) and pheatmap (version 1.0.12) R packages. Validation of division samples by senescence variable was conducted via subsetting normalized read counts by genes related to the Gene Ontology term “Cellular senescence” (GO 0090398). Differential expression analysis and log fold changes estimation were computed using DESeq2 for the last variable in the design formula controlling for cells senescence status, ouabain treatment reaction and an interaction of the indicated factors. To strengthen differential expression testing, log fold changes correction using combination of adaptive shrinkage estimator from the apeglm package (version 1.8.0) and specifying additional log fold change threshold equal to 0.667 was applied. Resulting shrunken estimates were used further for gene ranking and running Gene Set Enrichment Analysis using clusterProfiler (version 3.14.3) and fgsea (version 1.12.0) R packages with p-values adjustment for multiple comparisons according to the Benjamin-Hochberg method.

### Statistical analysis

To get significance in the difference between two groups Student’s t-test or Welch’s t-test was applied. For multiple comparisons between groups, ANOVA with Tukey HSD was used. Unless otherwise indicated, all quantitative data are shown as mean ± s.d., and the asterisks indicate significant differences as follows: ns, not significant, * p < 0.05, ** p < 0.01, *** p < 0.001. Statistical analysis was performed using R software.

## Acknowledgments

The authors are thankful to Maria Sirotkina for the assistance in the figures design.

## Author Contributions

A. V. B. supervised the work, wrote and edited the manuscript. A. V. B. and P. I. D. performed and designed most of the experiments. P. I. D. designed and conducted bioinformatic analysis, performed statistical analysis of the obtained data and edited the manuscript. A. N. S. performed FACS. I. I. M. determined intracellular ion contents. N. N. N. assisted in manuscript editing.

## Data Availability Statement

All data generated or analysed during this study are included in the manuscript and supporting files. Source data files have been provided for all Figures. Sequencing data have been deposited in GEO under accession codes GSE160702.

## Supplementary Information

**Figure 2–figure supplement 1.**
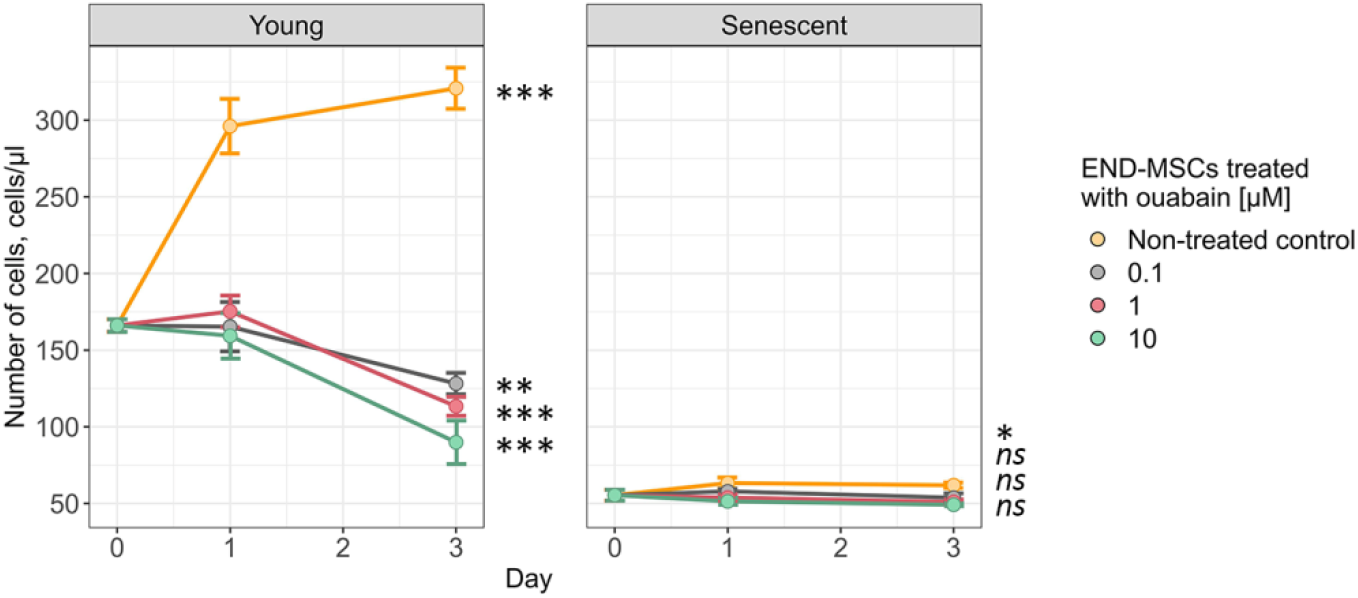
Standard growth curves of young and senescent END-MSCs treated with ouabain. Values are mean ± s.d. Statistical testing was performed using one-way ANOVA with Tukey HSD (results displayed are for Day 3 treatment outcomes against control at Day 0), n = 3, ns – not significant, ** p < 0.01, *** p < 0.001.

**Figure 4–figure supplement 1.**
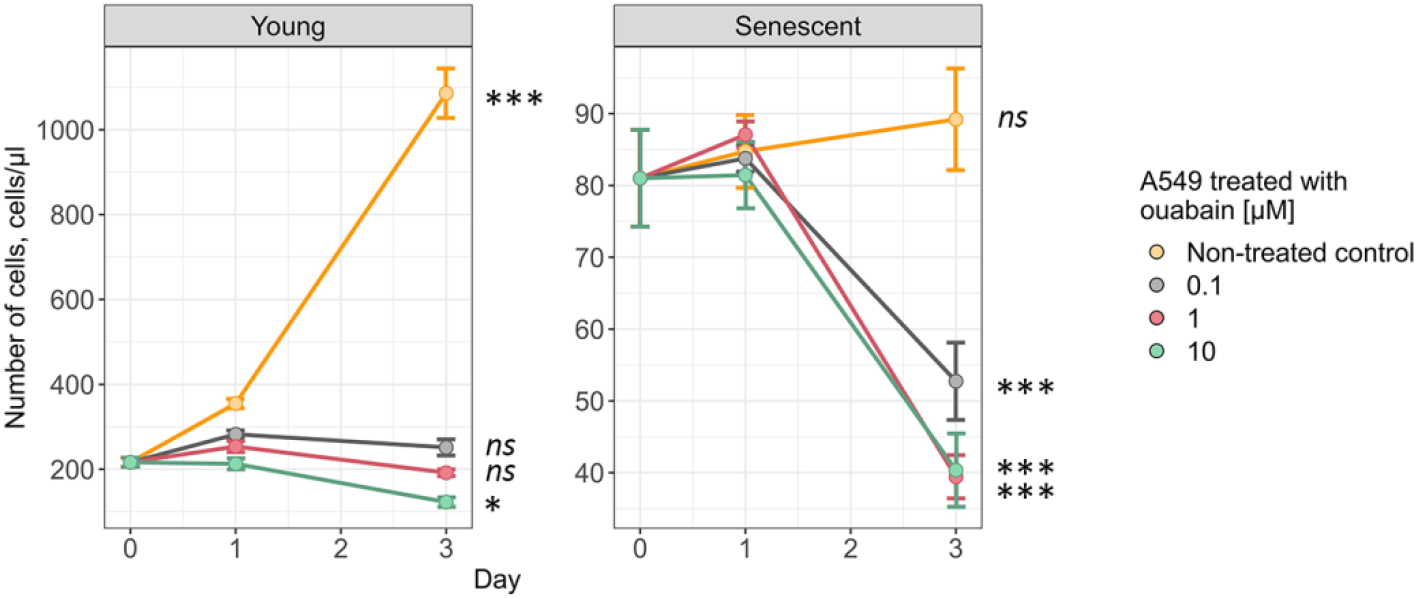
Growth curves of young and senescent A549 cells treated with ouabain. Values are mean ± s.d. Statistical testing was performed using one-way ANOVA with Tukey HSD (results displayed are for Day 3 treatment outcomes against control at Day 0), n = 3, ns – not significant, *** p < 0.001.

**Figure 5–figure supplement 1.**
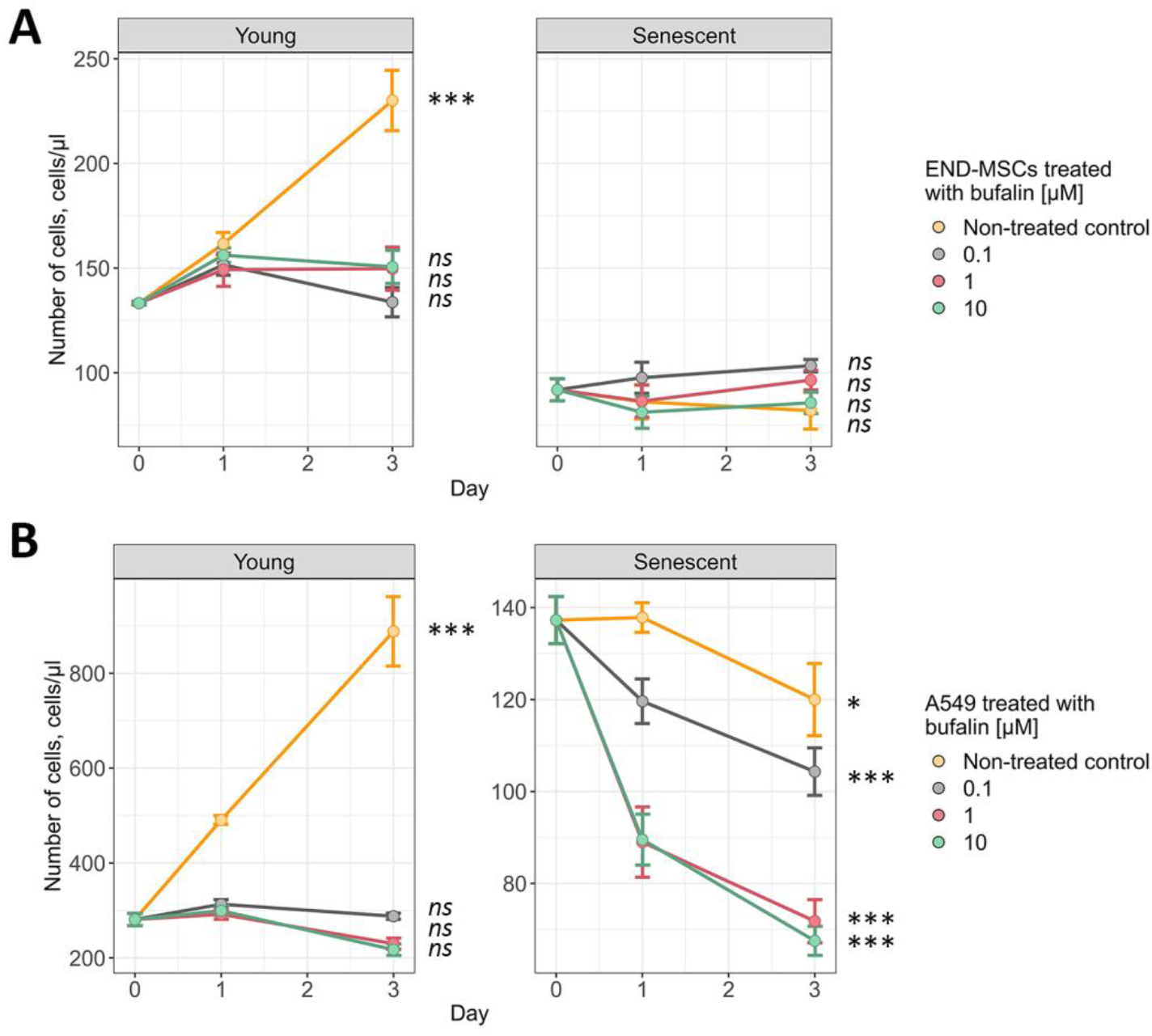
(**A**), (**B**) Standard growth curves of young and senescent END-MSCs and A549 treated with bufalin, respectively. Values are mean ± s.d. Statistical testing was performed using one-way ANOVA with Tukey HSD (results displayed are for Day 3 treatment outcomes against control at Day 0), n = 3, ns – not significant, * p < 0.05, *** p < 0.001.

**Figure 7–figure supplement 1.**
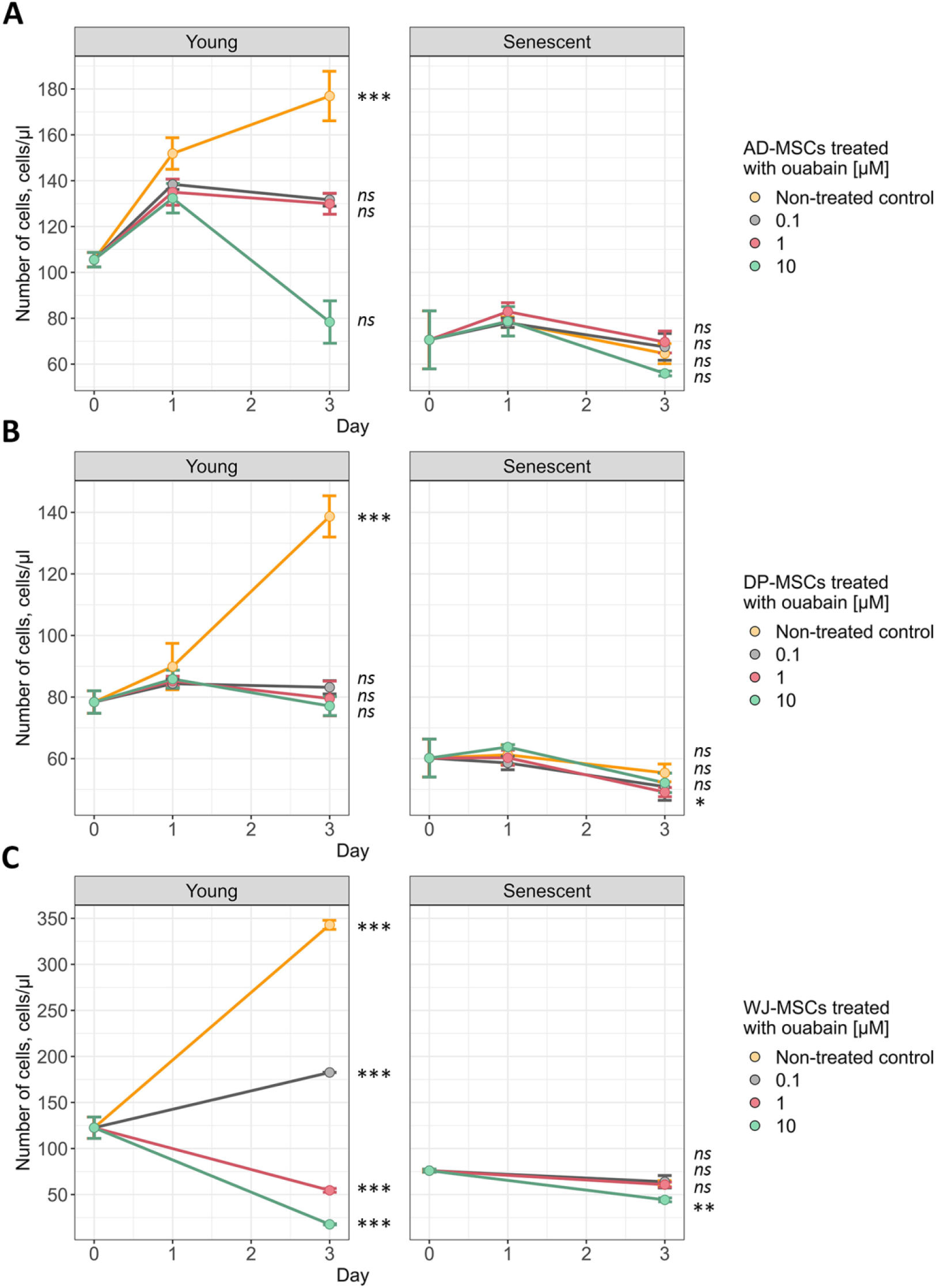
(**A**), (**B**), and (**C**) Standard growth curves of young and senescent AD-MSCs, DP-MSCs, and WJ-MSCs treated with ouabain, respectively. Values are mean ± s.d. Statistical testing was performed using one-way ANOVA with Tukey HSD (results displayed are for Day 3 treatment outcomes against control at Day 0), n = 3, ns – not significant, * p < 0.05, *** p < 0.001.

**Supplementary Figure 1.**
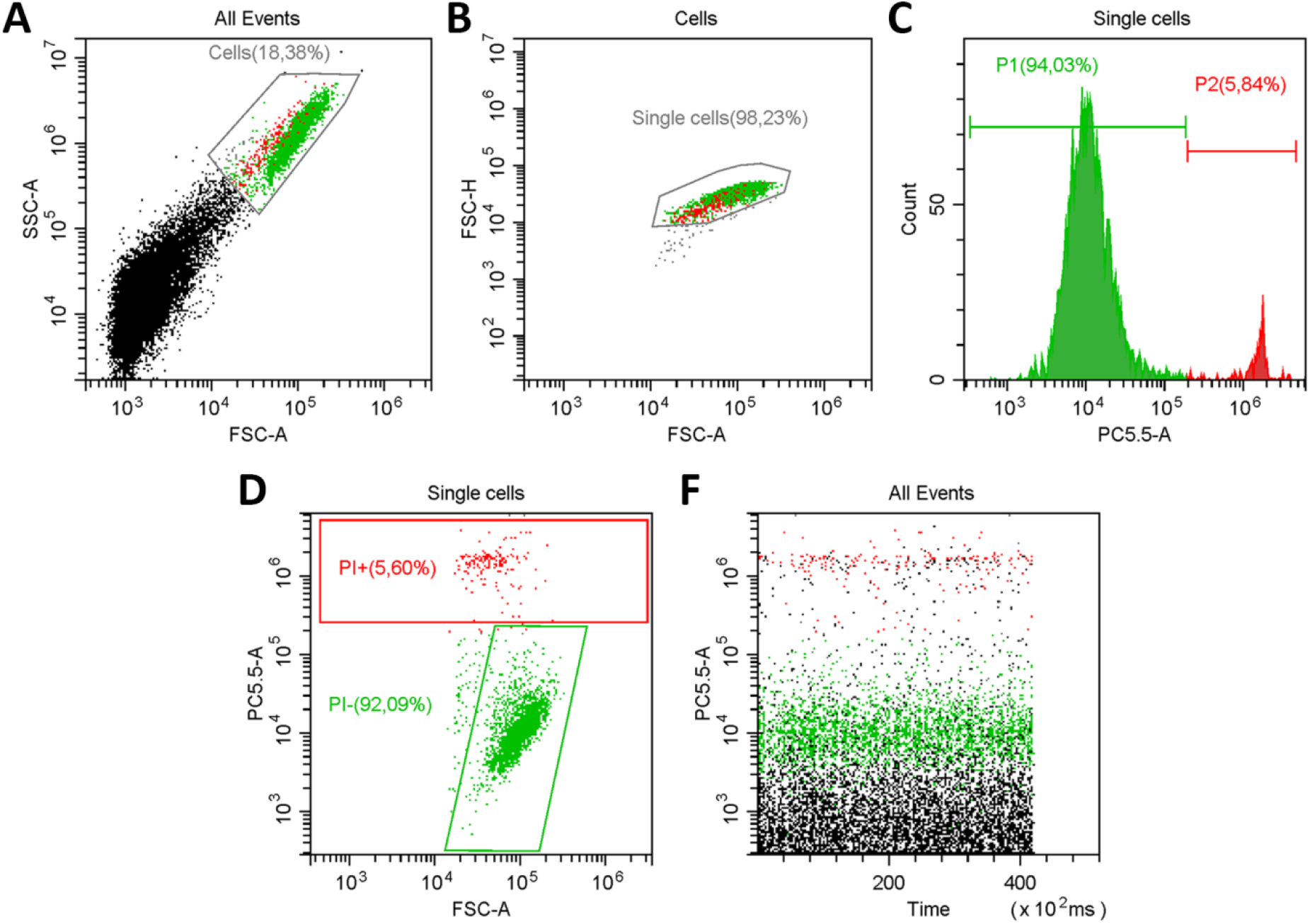
Flow cytometry gating strategy, related to the Figures of the main text. **a** Elimination of debris measurements identified as FSC-A low and SSC-A low. **b** Elimination of doublet measurements by gating along the region FSC-A/FSC-H. **c** One-parametric histogram reflecting viable and dead cells based on the PI (PC5.5-A) fluorescence intensity. **d** Dot plot reflecting viable and dead cells PI (PC5.5-A) fluorescence against FSC-A. **f** Time-resolution of sample flow.

**Supplementary Table 1**. The results of differential expression analysis for the cells resistant to ouabain-mediated senolysis (END-MSCs) against ones sensitive to ouabain-mediated senolysis (A549 and IMR-90).

**Supplementary Table 2**. Subset from the results of differential expression analysis for the cells resistant to ouabain-mediated senolysis (END-MSCs) against ones sensitive to ouabain-mediated senolysis (A549 and IMR-90) for genes encoding Na+/K+-ATPase subunits.

**Supplementary Table 3**. GSEA results for the differentially expressed genes between the cells resistant to ouabain-mediated senolysis (END-MSCs) and ones sensitive to ouabain-mediated senolysis (A549 and IMR-90) in the Gene Ontology Biological Processes terms.

